# Compensatory growth and recovery of tissue cytoarchitecture after transient cartilage-specific cell death in foetal mouse limbs

**DOI:** 10.1101/2023.06.20.545035

**Authors:** Chee Ho H’ng, Shanika L. Amarasinghe, Boya Zhang, Hojin Chang, David R. Powell, Alberto Rosello-Diez

## Abstract

A major question in developmental and regenerative biology is how organ size is controlled by progenitor cells. For example, while limb bones exhibit catch-up growth (recovery of a normal growth trajectory after transient developmental perturbation), it is unclear how this emerges from the behaviour of chondroprogenitors, the cells sustaining the cartilage anlagen that are progressively replaced by bone. Here we show that transient sparse cell death in the mouse foetal cartilage was repaired postnatally, via a two-step process. During injury, progression of chondroprogenitors towards more differentiated states was delayed, leading to altered cartilage cytoarchitecture and impaired bone growth. Then, once cell death was over, chondroprogenitor differentiation was accelerated and cartilage structure recovered, including partial rescue of bone growth. At the molecular level, ectopic activation of mTORC1 correlated with, and was necessary for, part of the recovery, revealing a specific candidate to be explored during normal growth and in future therapies.

## Introduction

Developmental robustness, i.e. the ability to achieve a relatively normal body plan despite genetic and environmental perturbations during development^2^, plays a key role in fitness and natural selection, but the underlying mechanisms are poorly understood. When it involves recovery of a normal growth trajectory after transient growth impairment, the phenomenon is referred to as catch-up growth^3^, which is related to–but distinct from–regeneration. In adult tissue regeneration, final organ mass had already been established before the injury, and there needs to be recovery of the <u>lost</u> mass. On the other hand, catch-up growth requires the recovery of <u>potential</u> mass, that is, the mass that would have been produced had the perturbation not happened. At the conceptual level, one potential scenario is that stem/progenitor cells can monitor a parameter directly or indirectly related to current organ size, and that this parameter dictates cell behaviours, but the specific molecular and cellular mechanisms remain obscure. It is also unclear why these recovery mechanisms are more powerful during foetal and perinatal stages than at juvenile stages, and the elucidation of this phenomenon will undoubtedly have a transformative impact for future regenerative therapies. Here, we addressed these fascinating questions using the vertebrate limb as a model, focusing on the long bones.

Growth of the long bones takes place by endochondral ossification, whereby a transient cartilage scaffold is progressively replaced by bone^4,5^. This replacement typically starts at the centre of the foetal skeletal elements, forming the primary ossification centre (Fig. 1a). In mice, secondary ossification centres form ∼1 week postnatally at both ends of the skeletal elements, so that, at each end, a cartilage disc (the growth plate) gets sandwiched between the two ossification centres. Cells in the growth plate (chondrocytes) traverse subsequent differentiation states from distal positions towards the bone centre: the resting zone (RZ) contains a pool of round quiescent progenitors, which in the proliferative zone (PZ) transition to flat proliferative chondrocytes stacked in longitudinal columns, and these cells then enlarge to form hypertrophic chondrocytes (HTCs, Fig. 1a). Many hypertrophic chondrocytes eventually die after laying down the extracellular matrix that will be replaced by bone matrix, laid down by osteoblasts. HTCs coordinate bone formation, vascularization, and cartilage resorption by attracting and communicating with the blood vessels that invade and degrade the cartilage from the surrounding tissues^6,7^.

**Fig. 1.**
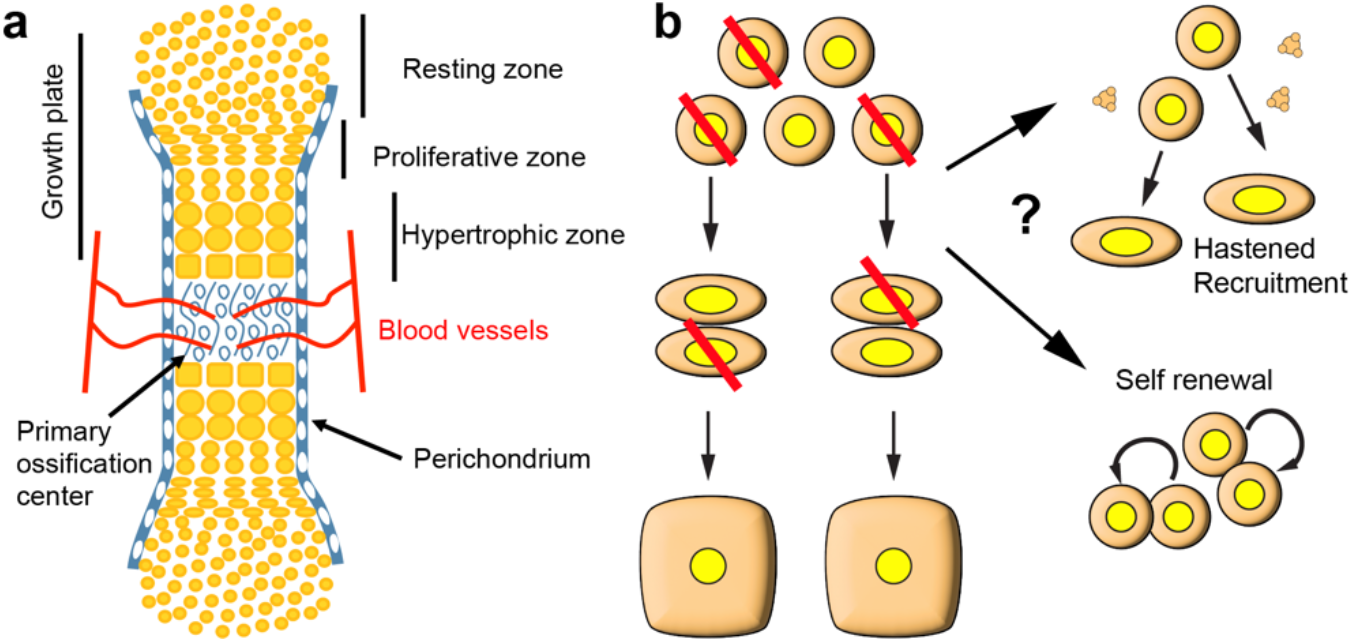
Model and approach to study foetal cartilage repair. **a** Schematic of a foetal skeletal element undergoing endochondral ossification. **b** Potential outcomes of mosaic cell ablation (red diagonal lines) in the foetal cartilage. Chondrocyte progenitors can either be immediately recruited into the proliferative population, using up their potential faster (eventually failing to catch-up). Alternatively, chondroprogenitors first self-renew, as shown by Oichi et al. ^1^, and then continue proliferating, leading to catch-up after a delay

The growth plate is regulated by multiple local and systemic signals^4,5^, and one of the key aspects that seems to be tightly controlled is the relative heights of the resting, proliferative and hypertrophic zones. Indeed, the proportion between the different growth-plate zones is a species-specific trait that correlates with bone size and growth rate, and that changes with age and anatomical location^8,9^. Yet, how this proportion is established is only partially understood. The size of the proliferative zone depends on a negative feedback loop between parathyroid hormone related peptide (PTHrP), produced by resting chondrocytes, and Indian hedgehog, produced by prehypertrophic ones (Fig. 1a). This feedback loop acts as a rheostat that couples proliferation and differentiation to maintain a certain growth plate size despite changes in cartilage production and degradation rates^10^. Regarding the cellular parameters affecting the actual growth rate, most of bone elongation has been shown to be contributed by the hypertrophic zone, via both cell enlargement and production of extracellular matrix^11^. Consequently, in growth plates whose HTCs reach very big sizes, changes in chondrocyte proliferation also exert a major effect on bone elongation, as they change the number of HTCs produced in a given time period^12^.

One approach to gain insight into the regulation of bone growth is to create structural defects in the cartilage and study the recovery (or lack thereof) of an adequate cartilage cytoarchitecture. Given that the mechanisms of foetal repair are more powerful than postnatal ones^13^, we posited that using this approach in foetal bones would be more informative than in postnatal ones. To gain insight into how bone growth rate is controlled, some labs (including ours) have studied the process of catch-up growth^2,3^. Three decades ago, Baron et al. showed that catch-up growth can happen after a local perturbation of bone growth^14^, suggesting that cartilage-intrinsic, and not systemic mechanisms, underlie the recovery. Subsequent studies refined this model, which became referred to as ‘autonomous’^15-18^. According to this model, chondrocyte progenitors have a limited intrinsic proliferative potential that is partially used up with every round of cell division until they become senescent. At this point the growth plate is consumed faster than it is replaced, and bone growth ceases. In this framework, catch-up growth is explained as follows: transient impairment of chondrocyte proliferation leads to chondrocyte progenitors preserving their proliferative potential, so that when the insult is lifted, their ‘age’ is lower than chronological age. Therefore, cartilage growth resumes at a speed associated with younger stages and lasts for a longer-than-normal period, until chondrocyte progenitors become senescent. This means that the growth trajectory is shifted horizontally towards a more advanced age, and the actual catch-up happens towards the normal end of the growth period. This model makes two strong predictions: 1) if the insult is mosaic (i.e., in a salt & pepper pattern), spared chondrocytes keep behaving normally and do not participate in the catch-up; 2) if chondrocytes are killed instead of arrested, there should not be catch-up, as the remaining cells keep using up their proliferative potential normally. The second prediction was tested in this study, as we shall show below. The first prediction was tested by one of us in a previous study, by inducing mosaic cell-cycle arrest in chondrocytes^19^. Unexpectedly, the spared chondrocytes showed enhanced proliferation rate, compensating for the proliferative arrest of the targeted ones. This cell-nonautonomous response was not compatible with a strict interpretation of the autonomous model and suggested that catch-up growth also involves cell-to-cell communication.

One of the cellular communication pathways that has been shown to impact bone growth is the insulin-like growth factor 1 (IGF1) pathway. While local IGF1 and IGF2 play a prominent role in the growth plate, only a minimum systemic threshold of liver-derived IGF1 is required for appropriate longitudinal bone growth ^20-22^. However, the local effects are region-specific. In resting cells, IGF1 promotes the recruitment towards proliferative chondrocytes, e.g. IGF1 receptor signalling represses PTHrP expression^23^. In addition, Oichi et al. showed that, during diet restriction in juvenile mice, *Axin2*^*+*^ progenitors in the cartilage were biased towards self-renewal instead of transitioning to the proliferative pool, which associated with decreased expression of *Igf1* in the resting zone^1^. Conversely, this bias and the expression of *Igf1* were reversed when the food supply was restored, leading to catch-up growth^1^. IGF1 also plays a key role in the hypertrophic zone (HZ). Indeed, IGF1 was shown to promote chondrocyte enlargement, alone or via interaction with other ligands^24-26^, and to mediate the differential enlargement exhibited by hypertrophic chondrocytes located in different growth plates of the same animal^26^.

One of the usual downstream effectors of IGF signalling that also targets resting and hypertrophic chondrocytes is the mechanistic target of rapamycin (mTOR, reviewed in ^27^). Specifically, it has been shown that mTOR complex 1 (mTORC1) promotes bone growth via stimulation of protein synthesis and cell size in the HZ^28-31^. On the other hand, postnatal activation of mTORC1 in the RZ associates with a switch to self-renewal and increased clonogenic capacity of chondrocyte progenitors^32^. In this regard, mTORC1 was shown to enhance expression of PTHrP^33^, a marker of so-called cartilage stem cells^34^. In summary, since mTORC1 regulates growth-related processes across multiple regions of the growth plate, it stands as a main candidate to coordinate the relative height of growth plate zones, and associated bone growth parameters.

In this study, we induced mosaic cartilage damage to determine whether and how cartilage is repaired afterwards. To test the second prediction of the autonomous model of catch-up growth, we investigated the response to transient mosaic cell death induced in the cartilage of the left foetal long bones (Fig. 1b), with the right limb as internal control. We predicted two potential scenarios at the level of resting chondrocytes. In one scenario, the remaining chondroprogenitors accelerate their recruitment towards the proliferative pool, but some are used to fill-in the gaps left by the dying ones, instead of being used to promote elongation, leading to a progressively increasing length difference with the control limb (Fig. 1b, top). In the alternative scenario, the injury could induce self-renewal of some of the spared chondroprogenitors at the expense of the PZ pool, leading to an initial growth delay, followed by elongation at the rate of a younger animal, and catch-up after a longer time (Fig. 1b bottom). The latter scenario was recently found in the case of catch-up growth induced by transient diet restriction^1^. In contrast, we unexpectedly found that a hybrid compensatory growth took place via two mechanisms: 1) replenishment of resting chondrocytes due to a transient bias towards self-renewal, followed by accelerated transition to the proliferative pool, but insufficient to gain back the lost size; 2) transiently increased hypertrophic chondrocyte size. These responses were associated with increased phosphorylated ribosomal protein S6 (p-S6) signal, partially dependent on mTORC1 signalling. Moreover, the compensatory response led to an almost perfect recovery of the zone proportions in the injured growth plates, a process which was impaired by inhibition of mTORC1 signalling, in line with the modulatory effect that p-S6 has been recently shown to play in skin wound healing^35^. These results suggest that the regulative behaviour of chondroprogenitors is more complex than expected, and reveal new molecular candidates to target in order to improve or extend the reparative phase.

## Materials and Methods

### Animal models

The use of animals in this study was approved by the Monash Animal Research Platform animal ethics committee at Monash University. The *Pitx2-ASE-Cre* (aka *Pitx2-Cre*) mouse line, obtained from Prof. Hamada^36^, was crossed with the *Col2a1-rtTA* mouse line^37^, and then inter-crossed to generate *Pitx2-Cre/Cre; Col2a1-rtTA* mice. Genotyping was performed as described previously^19^. The *Tigre*^*Dragon-DTA*^ mouse line was described in ^38^. Experimental and control animals were generated by crossing *Pitx2-Cre/Cre; Col2a1-rtTA/+* females with *Tigre*^*Dragon-DTA/DTA*^ males (i.e. homozygous for the conditional misexpression allele). The separation of control and experimental animals was based on the rtTA genotype. Pregnant females were administered doxycycline hyclate (Sigma, 1 mg/ml in drinking water, with 0.5% sucrose for palatability) from E12.25 to E13.75. The day of vaginal plug detection was designated as E0.5, and E19.5 was referred to as P0.

### Drug administration

A solution of Rapamycin (Selleckchem #S1039) was prepared at the concentration of 2 mg/ml in a mixture of water, 2% dimethylsulfoxide (DMSO), 30% polyethylene glycol 300 (PEG 300), and 5% polysorbate 80 (Tween 80). This solution was administered to the pregnant female via intraperitoneal (i.p.) injection at a dose of 4 mg/kg (twice daily).

### EdU incorporation and detection

A solution of 5-Ethynyl-2’-deoxyuridine (EdU) was prepared at 6 mg/ml in phosphate-buffered saline (PBS). This solution was administered at a dose of 30 µg/g, subcutaneously (s.c) for pups and intraperitoneally (i.p.) for pregnant females, 1.5 hours before euthanizing the mice (or 2 days before, for pulse-chase experiments). To detect EdU, a click chemistry reaction with fluorescein-conjugated azide (Lumiprobe #A4130) was performed once the immunohistochemistry and/or TUNEL staining were completed on the same slides. Briefly, the working solution was prepared in PBS, adding CuSO_4_ (Sigma # C1297) to 4 mM, the azide to 0.4 µM and freshly-dissolved ascorbic acid (Sigma #A0278) to 20 mg/ml, and incubating the sections for 15 min at room temperature in the dark, followed by PBS washes.

### Sample collection and processing

Mouse embryos were euthanized using hypothermia in cold PBS, while mouse pups were euthanized by decapitation. Upon collection of the embryos or pups, the limbs (including full tibiae and/or femora) were carefully dissected out in cold PBS, skinned, and fixed in 4% paraformaldehyde (PFA) for 2 days at 4°C. Samples of P1 or younger were not subjected to decalcification. For P3, P5, and P7 samples, decalcification was performed by immersing the specimens in 0.45M ethylenediaminetetraacetic acid (EDTA) in PBS for 3, 5, and 7 days, respectively, at 4°C. Following several washes with PBS, the limb tissues were cryoprotected in PBS containing 15% sucrose and then equilibrated in 30% sucrose at 4°C until they sank. The hindlimbs were then oriented sagittally, facing each other, with the tibiae positioned at the bottom of the block (closest to the blade during sectioning) and embedded in Optimal Cutting Temperature (OCT) compound using cryomolds (Tissue-Tek). The specimens were frozen by immersing them in dry-ice-cold iso-pentane (Sigma). Serial sections with a thickness of 7 µm were collected using a Leica Cryostat on Superfrost slides. The sections were allowed to dry for at least 30 minutes and stored at -80°C until further use. Prior to conducting histological techniques, the frozen slides were brought to room temperature in PBS, and the OCT compound was washed away with additional rounds of PBS rinses.

### Micro-CT and bone length analysis

The micro-CT and bone length analysis was as previously described^39^. Briefly, samples were retained and fixed in 4% PFA as residual tissues from other experiments in the Rosello-Diez lab. Whole femora and humeri were scanned using a Siemens Inveon PET-SPECT-CT small animal scanner in CT modality (Monash Biomedical Imaging). The scanning parameters included a resolution of 20 and 40 µm, 360 projections at 80 kV, 500 µA, 600 ms exposure with a 500 ms settling time between projections. Binning was applied to adjust the resolution with 2 × 2 for 20 µm scans and 4 × 4 for 40 µm scans. The acquired data were reconstructed using a Feldkamp algorithm and converted to DICOM files using Siemens software. For the analysis and bone length measurements, Mimics Research software (v21.0; Materialize, Leuven, Belgium) equipped with the scripting module was utilized to develop the analysis pipeline.

### Histology, Haematoxylin and Eosin and Alcian Blue staining

Following sectioning, the sections were washed in distilled water. For Alcian blue staining, the sections were incubated in a 1% Alcian Blue solution in distilled water, adjusted to pH 1.0 with hydrochloric acid (Sigma #A5268), for 15 minutes at room temperature, followed by rinsing in water. For Haematoxylin and Eosin (H&E) staining, the sections were stained with filtered mercury-free Harris haematoxylin solution (Point of Care Diagnostics #VWRC351945S) for 5 minutes, washed in running tap water for 5 minutes, and then rinsed in 95% ethanol. Next, the sections were counterstained with a 0.25% Eosin Y solution (Sigma #HT110116) in 80% ethanol and 0.5% glacial acetic acid for 1 minute. For both Alcian Blue and H&E staining, dehydration was performed by passing the sections through 95% ethanol, absolute ethanol, and xylene. Finally, the sections were mounted using a xylene-based mounting medium, dibutylphthalane polystyrene xylene (DPX) (Sigma #100579).

### Immunohistochemistry and TUNEL staining

For antigen retrieval, the sections were subjected to citrate buffer (10 mM citric acid, 0.05% Tween 20 [pH 6.0]) at 90°C for 15 minutes. Afterward, the sections were cooled down in an ice water bath, washed with PBSTx (PBS containing 0.1% Triton X-100). To perform TUNEL staining, the endogenous biotin was blocked using the Avidin/Biotin blocking kit (Vector #SP-2001) after antigen retrieval. Subsequently, TdT enzyme and Biotin-16-dUTP (Sigma #3333566001 and #11093070910) were used according to the manufacturer’s instructions. Biotin-tagged DNA nicks were visualized using Alexa488-or Alexa647-conjugated streptavidin (Molecular Probes, diluted 1/1000) during the incubation with the secondary antibody.

For immunohistochemistry staining, sections were incubated with the primary antibodies prepared in PBS for either 1.5 hours at room temperature or overnight at 4°C (see list of antibodies below). Following PBSTx washes, the sections were incubated with Alexa488-, Alexa555-, and/or Alexa647-conjugated secondary antibodies (Molecular Probes; diluted 1/500 in PBSTx with DAPI) for 1 hour at room temperature. After additional PBSTx washes, the slides were mounted using Fluoromount™ Aqueous Mounting Medium (Sigma). The antibodies used, along with their host species, vendors, catalogue numbers, and dilutions, were as follows: mCherry (goat polyclonal, Origene Technologies #AB0040-200, diluted 1/500), p-S6 (rabbit polyclonal, Cell Signaling Technology #2211S, diluted 1/300).

### In situ hybridization

To obtain the DTA riboprobe, the coding sequence of DTA was first amplified from the pB6-Rosa26-DTA-Soriano-for vector using primers XhoI_Kozak_DTA_F (gactgacctcgaggccaccATGGAAGCGGGTAGGCCTT) and ClaI_DTA_R (gactgacatcgatTTAGAGCTTTAAATCTCTGTAGGTAGTT), and directionally cloned into XhoI-ClaI digested pBS KS plasmid. The resulting plasmid was linearized with XhoI and purified by phenol/chloroform extraction. In vitro transcription was performed on the linearized template using T7 RNA polymerase (Roche DIG-labelling mix) to obtain a full-length Digoxigenin-labelled *DTA* riboprobe, which was purified via the LiCl/EtOH precipitation method, and resuspended in RNAse-free water to a concentration of ∼400 ng/µl. The pB6-ROSA26-DTA-Soriano-for vector was a gift from Mario Capecchi (Addgene plasmid # 125745 ; http://n2t.net/addgene:125745 ; RRID:Addgene_125745).

To perform *in situ* hybridisation, sections were fixed in 4% PFA for 20 minutes at room temperature, washed in PBS, and treated with 4µg/ml Proteinase K for 15 minutes at 37°C. After washing in PBS, the sections were refixed with 4% PFA, followed by treatment with 0.1N pH8 triethanolamine (Sigma #90279), 0.25% acetic anhydride (Sigma #320102) for 10 minutes at room temperature. Subsequently, the sections were washed in PBS and water and incubated with prehybridization buffer (50% formamide, 5x SSC pH 5.5, 0.1% 3-[(3-Cholamidopropyl)dimethylammonio]-1-propanesulfonate (CHAPS), 0.05 mg/ml yeast tRNA, 0.1% Tween 20, 1x Denhardt’s) at 60°C for 30 minutes. The sections were then incubated with 1µg/ml preheated riboprobes and subjected to hybridization at 60°C for 2 hours. Post-hybridization washes were performed using post-hybridisation buffer I (50% formamide, 5x pH 5.5 SSC, 1% SDS) and II (50% formamide, 2x pH 5.5 SSC, 0.2% SDS) preheated at 60°C for 30 minutes, respectively. The sections were then washed with maleic acid buffer (MABT: 100mM maleic acid, 150mM NaCl, 70mM NaOH, 0.1% Tween 20) and blocked with 10% goat serum, 1% blocking reagent (Roche #11096176001) in MABT at room temperature for 30 minutes. Next, the sections were incubated overnight at 4°C with anti-digoxigenin-AP (Sigma #11093274910) diluted 1/4000 in MABT with 2% goat serum and 1% blocking reagent. After several MABT washes, the sections underwent incubation with AP buffer (0.1M Tris-HCL pH 9.5, 0.1M NaCl, 0.05M MgCl_2_, 0.1% Tween 20), with the second one containing 1mM levamisole and then developed colour using BM purple (Roche #11442074001) at 37°C. Following a wash in PBS, the sections were fixed in 4% PFA, counterstained with Nuclear Fast Red (Sigma #N3020) at room temperature for 10 minutes, and rinsed in water. Dehydration was performed by passing the sections through 70%, 90% ethanol, absolute ethanol, and xylene. Finally, the sections were mounted with DPX (Sigma #100579).

### Imaging

Sagittal sections of the limbs were captured, with a focus on the area between the distal femora and proximal tibiae. Typically, at least 2 sections per limb were analyzed, although in most cases 4 sections were examined. In the case of cultured distal femora, frontal sections were used as they provided better identification of the different epiphyseal regions. To determine the boundaries of the resting zone (RZ), proliferative zone (PZ) and hypertrophic zone (HZ), morphological criteria were applied. The transition between round (resting) and flat (columnar) nuclei, forming an arch along the upper point of the grooves of Ranvier, was considered as the start of the PZ. On the other hand, the transition towards larger, more spaced nuclei (pre-hypertrophic) marked the end of the PZ. The point where the pericellular matrix exhibited a sharp reduction around enlarging chondrocytes was designated as the beginning of the HZ. The distal end of the last intact chondrocyte served as the endpoint of the HZ. For imaging, bright-field and fluorescence images were acquired using a Zeiss inverted microscope (Imager.Z1 or Z2) equipped with Axiovision software (Zeiss/ZenBlue). Mosaic pictures were automatically generated by assembling individual tiles captured at 10× magnification for bright-field images or 20× magnification for fluorescence images.

### Image analysis and quantification

The regions of interest such as resting, proliferative and hypertrophic zones (RZ, PZ, HZ) and the secondary ossification centre (SOC) were identified from imaged sections of the multiple models (left and right; experimental and control proximal tibial cartilage). Consistent parameters such as brightness, contrast, filters and layers were kept the same for all the images in the same study.

### Cell count analysis

The RZ and PZ were identified and segmented from sections stained for DAPI, tdTomato, and TUNEL or EdU using FIJI software. The number of cells in the region of interest (RZ & PZ) was measured in using Cell Profiler.

### Cell size analysis

The HZ was identified and segmented from sections stained for Alcian Blue. The area (µm^2^) of individual hypertrophic chondrocyte was measured using FIJI software. Data from at least 3 mice (6 limbs), and 2-4 sections per limb, was pooled together to build the histograms in Graphpad Prism.

### Cartilage length analysis

Using H&E staining, distinct zones of cartilage were identified. The length (µm) of each cartilage zone (RZ, PZ, HZ) was then measured using FIJI software.

### p-S6 area percentage analysis

Following DAPI and p-S6 staining, the area percentage (%) of p-S6 signals over the total area of region of interest (RZ + PZ) were measure based on the signal threshold using FIJI software.

### Secondary Ossification Centre (SOC) relative area analysis

Following H&E staining, the SOC area (including blood vessel invasion areas and hypertrophic cells) and the total epiphyseal area were measured using FIJI software.

### Statistical analysis

Statistical comparisons were performed using appropriate tests based on the experimental design. An unpaired t test was used for comparisons involving one variable and two conditions. Two-way ANOVA was utilized for comparisons involving two variables and two or more conditions. Non-parametric tests were selected when the assumption of normality could not be met. All statistical analyses were conducted using Prism9 software (GraphPad).

### Multiplexed RNA-seq and analysis

#### Experimental Design

The RNA-Seq study has a 2 (genotype: control, experimental) by 2 (side: left, right) by 4 (stage: E15.5, E17.5, P0, P3) factorial design. We ensured a minimum number of 3 biological replicates per experimental group were sequenced (2 for controls). The final experimental design is summarised in Extended Data Fig. 7. For the stages P0 and P3 there were two technical replicates per biological sample, which were subsequently merged after confirming their similarity.

#### RNA extraction, multiplexed library preparation, sequencing

Control and experimental samples (2-3 biological replicates each) were collected for 4 different stages: E15.5, E17.5, P0 and P3. Left and right cartilage samples (each containing proximal and distal tibia and femur) were kept separated for each sample, and flash-frozen upon dissection. Total RNA was extracted using the Monarch Total RNA Miniprep Kit (New England Biolabs), following manufacturer instructions. From this, RNA-Seq of messenger RNAs (mRNAs) was performed using a custom in-house multiplex method, which allowed us to sequence up to 24 different samples in the same sequencing lane. Briefly, samples were given a unique i7 index (together with UMI) during individual polyA-priming and first-strand synthesis, which also added a template-switch sequence to the 5’ end. Samples were then pooled and amplified using P7 and an oligo which binds the template-switch sequence. Final library construction was completed by tagmentation and addition of P5 (with i5 index) by PCR. Sequencing was performed on an Illumina NSQ2k run with up to 101nt SR (cDNA). An 18-nt i7 read contains the 8-nt index and 10-nt UMI and, where required, an 8-nt i5 index read is also generated. We obtained a read depth of at least 15 million reads per library.

#### RNA-Seq data analysis

Raw data was examined by the program FASTQC and MULTIQC for read quality, detection of adapter contaminations and presence of overrepresented sequences^40^. i7 index reads were trimmed (the first 8bp removed) so the remaining 10-bp UMI can be analysed and data filtered per sample before mapping using cutadapt programme^41^. The tools umi-tools^42^ (v 0.5.5) and Bbmap^43^ (v. 38.81) were used to demultiplex the samples based on barcodes from the multiplexed data. Next, STAR aligner^44^ (v 2.7.9a) was used to map the fastq reads to the mm39 (GRCm39) reference genome from Ensembl with the following parameters:--outFilterMultimapNmax 20, --outFilterMismatchNmax 5, --outFilterScoreMinOverLread 0. featureCounts() function from the R/Bioconductor package Rsubread^45^ (v 2.10.4) was used to generate the gene count matrix for the aligned reads with following parameters; allowMultiOverlap = TRUE, minOverlap = 1, fracOverlap = 0, fracOverlapFeature = 0.

LIMMA (Linear Models for Microarray Data) package (v 3.52.4) implemented in the R software environment ^46,47^ (v 4.2.2) was used to calculate the t-statistics for mean expression values for each gene. The four stages were analysed separately. Surrogate variables were included in the design matrices to further remove any unwanted variations^48^. The read count data fed into the LIMMA linear model fitting were transformed using Voom with quantile normalization^49^ followed by group-means parameterization and robust eBayes^47^. The contrast matrix was created with the final goal of identifying the differentially expressed genes in limb that is due to the definite effect of the perturbed growth within each stage. p-values were corrected for multiple testing according to Benjamini and Hochberg (1995) to control the false discovery rate (FDR)^50^. Genes were considered significantly different in expression relative to control if the FDR-adjusted p-value ≤ 0.1.

We analysed the expression of canonical molecular signatures from MSigDB^51^ using ROAST test from the edgeR package^52^ (v 3.38.4), a self-contained, rotation gene set testing which tests whether the genes within a gene set are differentially expressed. The plots were created using ggplot2 package^53^ and pheatmap^54^ packages from R/Bioconductor and the *Degust* online tool^55^.

Weighted gene co-expression analysis was done within each stage using the R package WGCNA^56^. The gradient method was used to test the independence and the average connectivity degree of different modules with different power value (the power value ranging from 1 to 20). The appropriate power values for the four stages were established when the degree of independence was highest. Once the power value was identified, the module construction was conducted through the manual network construction approach of WGCNA. The minimum number of genes per cluster was set as 30 for the high reliability of the results.

Functional annotation of identified genes from both gene expression and gene co-expression analyses was done using web applications such as DAVID^57^, Panther Pathway^58^ and STRING^59^.

## Results

### A model of acute unilateral cell death in the foetal limb cartilage

To induce transient cell death in the cartilage of the left developing limbs, we utilized a double conditional mouse strain (*Tigre*^*Dragon-DTA*^, *Dragon-DTA* hereafter) that we previously generated^38^, in which expression of attenuated diphtheria toxin fragment A (DTA) requires previous or current Cre activity as well as active tetracycline-responsive transactivator (tTA) or its reverse version rtTA. Cre activity is required to prime DTA expression by removing a floxed tdTom-STOP cassette, and (r)tTA drives transcription of either tdTomato (in the unrecombined allele) or DTA (in the recombined one) from a Tet-responsive element (TRE) located upstream of the transgene (Fig. 2A). Importantly, the activity of (r)tTA can be controlled by the Tetracycline analogue Doxycycline (Dox, which activates rtTA and inhibits tTA), such that the model is inducible and reversible^38^. We crossed males homozygous for this allele with females homozygous for *Pitx2-ASE-Cre*^36^ (*Pitx2-Cre* hereafter) and hemizygous for *Col2a1-rtTA*^37^, to generate the *Pit-Col-DTA* model. The *Pitx2-Cre* allele drives Cre expression in the left lateral plate mesoderm (which gives rise to the limb mesenchyme), while *Col2a1-rtTA* drives rtTA expression in the *Col2a1-*expressing chondrocytes^37^, so that their activities intersect in the cartilage of the left limbs, killing chondrocytes in those skeletal elements^38^. To achieve a brief peak of cell death, but intense enough to generate an obvious left-right asymmetry, we first tested several Dox regimes in the drinking water, varying the concentration and duration of treatment. 0.5 mg/ml Dox given from embryonic day (E) 12.5 to E15.5 caused limb asymmetry in some specimens, but this result was not reproducible unless Dox concentration was increased to 1 mg/ml (not shown). When 1 mg/ml Dox was given for 12h and the embryos collected right after, incipient tdTomato signal was detected in the right cartilage, indicating that the TRE had been active for a few hours, although cell death (assessed via TUNEL staining) was not detected yet (Ext. Data Fig. 1a). Interestingly, when Dox was given from E12.25 and the embryos collected 30h later (at E13.5), we detected clear expression of *DTA* in the left cartilage, but still not cell death (Ext.

**Fig. 2.**
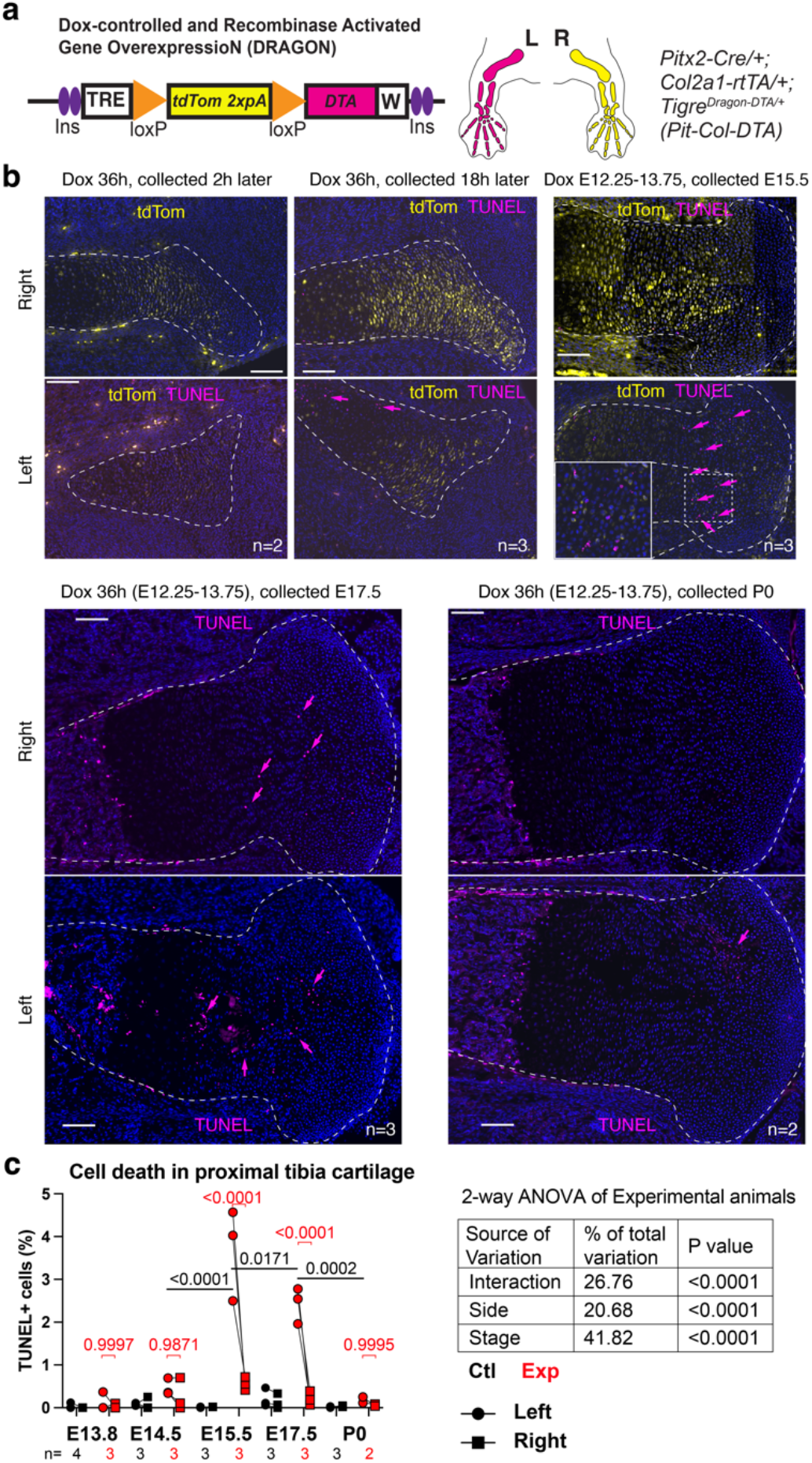
A model of transient unilateral cell death in the foetal cartilage. **a** Genetic combination to achieve unilateral, cartilage-specific, inducible and reversible DTA expression. **b** Examples of the tdTomato expression and cell death (TUNEL, arrows) achieved in the cartilage (dashed lines) after the indicated Dox treatments. **c** Quantification of TUNEL^**+**^ cells at the indicated stages/ locations. 2-way ANOVA table is shown, with the posthoc multiple comparison tests (across sides and across stages in Exp cartilage) shown above the graphs.

Data Fig. 1b, see also Fig. 6c). Similarly, when Dox was given from E12.25 to E13.75 (i.e., for 36h) and the embryos collected 2-4h later (E13.8), clear tdTomato but little TUNEL signals were detected (Fig. 2b, c). After a similar 36-h Dox pulse, the first clear indication of cell death in the left cartilage was found 18h later (∼E14.5), and especially expanded at 42h post-Dox withdrawal (∼E15.5, Fig. 2b, c). The levels of cell death were still quite high 2 days later, at E17.5, but almost completely gone by E19.5, i.e. postnatal day 0 (P0, Fig. 2b, c). The results thus show that transient Dox treatment in the *Pit-Col-DTA* model indeed triggers acute cell death. Of note, the right experimental limb (i.e. the internal control) showed a few DTA^+^ cells at E13.5 (Ext. Data Fig. 1b, arrowheads) and a trend towards increased cell death as compared to absolute controls at E15.5 (Fig. 2c). This was somewhat expected, as AR-D showed that *Pitx2-Cre* exhibits some minor activity in the right limb too^19^.

### Cartilage is almost completely repaired 1 week after injury

After confirming the transient peak of cell death in the *Pit-Col-DTA* model, we analysed the effect on cartilage integrity and cytoarchitecture. Haematoxylin and eosin staining of the proximal growth plate at multiple stages showed obvious “gaps” in the E17.5-E18.5 left experimental cartilage (Exp L), spreading across the resting and proliferative zones (Fig. 3a and not shown). By P0, the damage had expanded partially or completely to the HZ, showing big acellular patches, filled only with extracellular matrix (Fig. 3b). Most remarkably, however, by P3-P5 most of the damage in the resting zone had healed, and the gaps in other zones were less obvious in most cases (Fig. 3c, d. Only 1 out of 6 Exp L tibia showed empty patches bigger than 4 cell diameters). By P7, no more cartilage gaps were detected (Ext. Data Fig. 2a). However, some long-term effects of the acute injury were still detectable at P7. For example, the whole cartilage was slightly shorter in the Exp L tibia as compared to controls, although no differences between Exp L and Exp R were found (Fig. 3e). Lastly, we noticed that formation of the secondary ossification centre (SOC) was delayed in the P7 Exp L proximal tibia, suggesting that the cartilage repair process entails an overall delay of the endochondral ossification process (Ext. Data Fig. 2b).

**Fig. 3.**
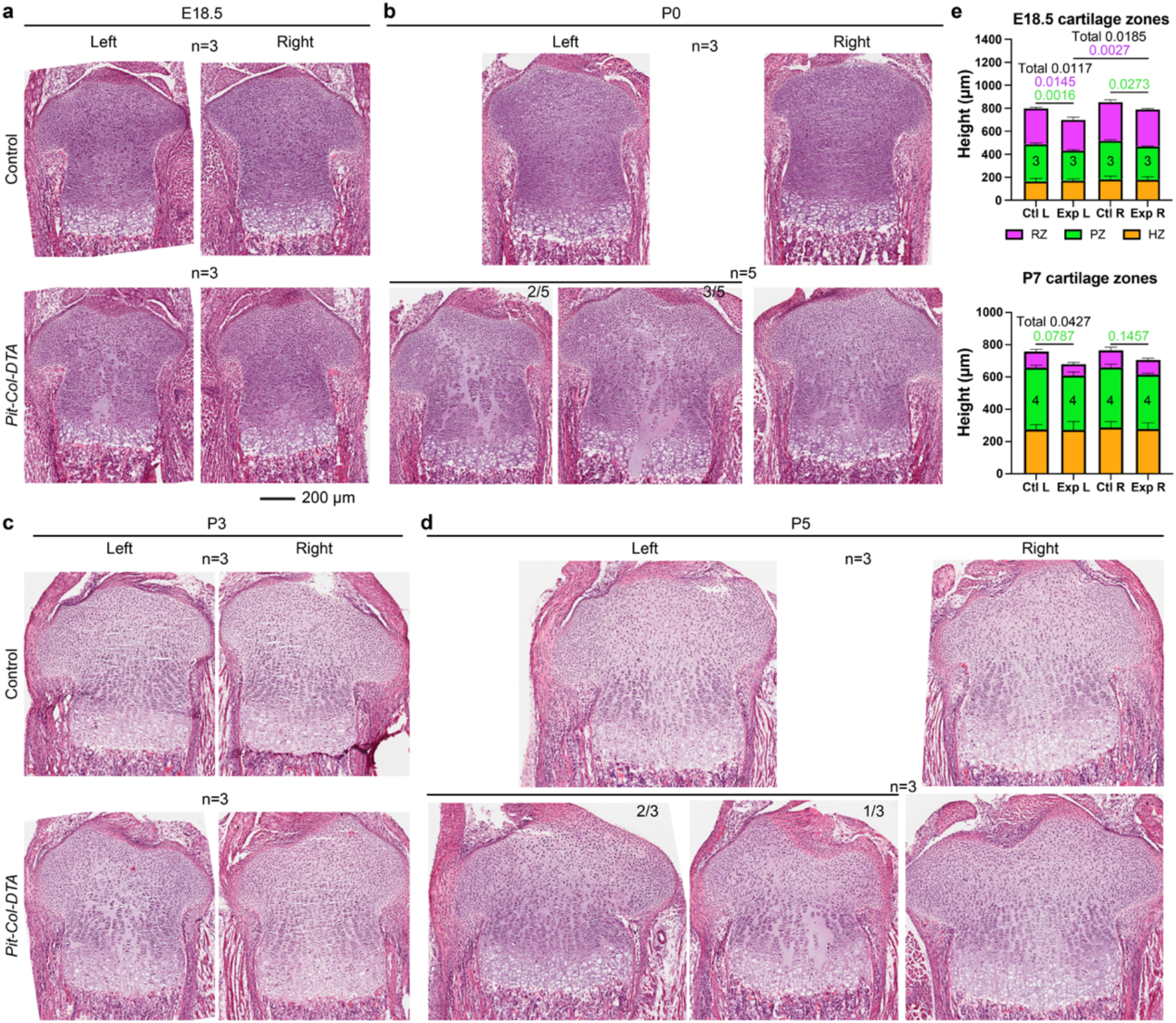
After transient cell death, cartilage cytoarchitecture is repaired within 1 week. **a-d** H&E staining on sections of the proximal tibia at the indicated embryonic (E) and postnatal (P) stages. **e** Quantification of the maximum length of each cartilage zone (RZ/PZ/HZ, resting/proliferative/hypertrophic) at the indicated stages, categorized by genotype and side. Number of samples as indicated inside the green zone of the bar graphs. Statistical analyses are 2-way ANOVAs and Sidak’s post-hoc multiple comparisons tests, performed at each stage for all zones at the same time, in a pairwise manner (CtlL-ExpL, CtlR-ExpR, CtlR-ExpR, ExpL-ExpR).

### Transient unilateral cell death in the developing limb cartilage leads to limb asymmetry, followed by relative catch-up growth

We next analysed the effect of unilateral transient cartilage cell-death on long-bone length and symmetry, using whole-body micro-CT and semi-automatic length analysis, as we described ^39^. At E17-P0, when cell death is past its peak, the left/right ratio of bone length reaches an average value of 0.77 for the femur, 0.90 for the humerus (Fig. 4a, b and Ext. Data Fig. 3). The difference between forelimb and hindlimb is consistent with the fact that the *Pitx2-Cre* allele is more active in the prospective hindlimb region than the forelimb one^19^. Interestingly, by P2-3 the left/right femur ratio was significantly increased to 0.91 (Fig. 4a and Ext. Data Fig. 3), indicating partial recovery of relative symmetry. This ratio plateaued at 0.9-0.95 by P5-P7, not showing further recovery by P14 (Fig. 4a), and keeping a similar value at P100, well past the end of the normal growth period (Fig. 4a and Ext. Data Fig. 3). The results thus show that, against the prediction of the autonomous model, cell death in the cartilage triggers compensation, which can only be cell-nonautonomous (i.e. driven by the spared cells).

**Fig. 4.**
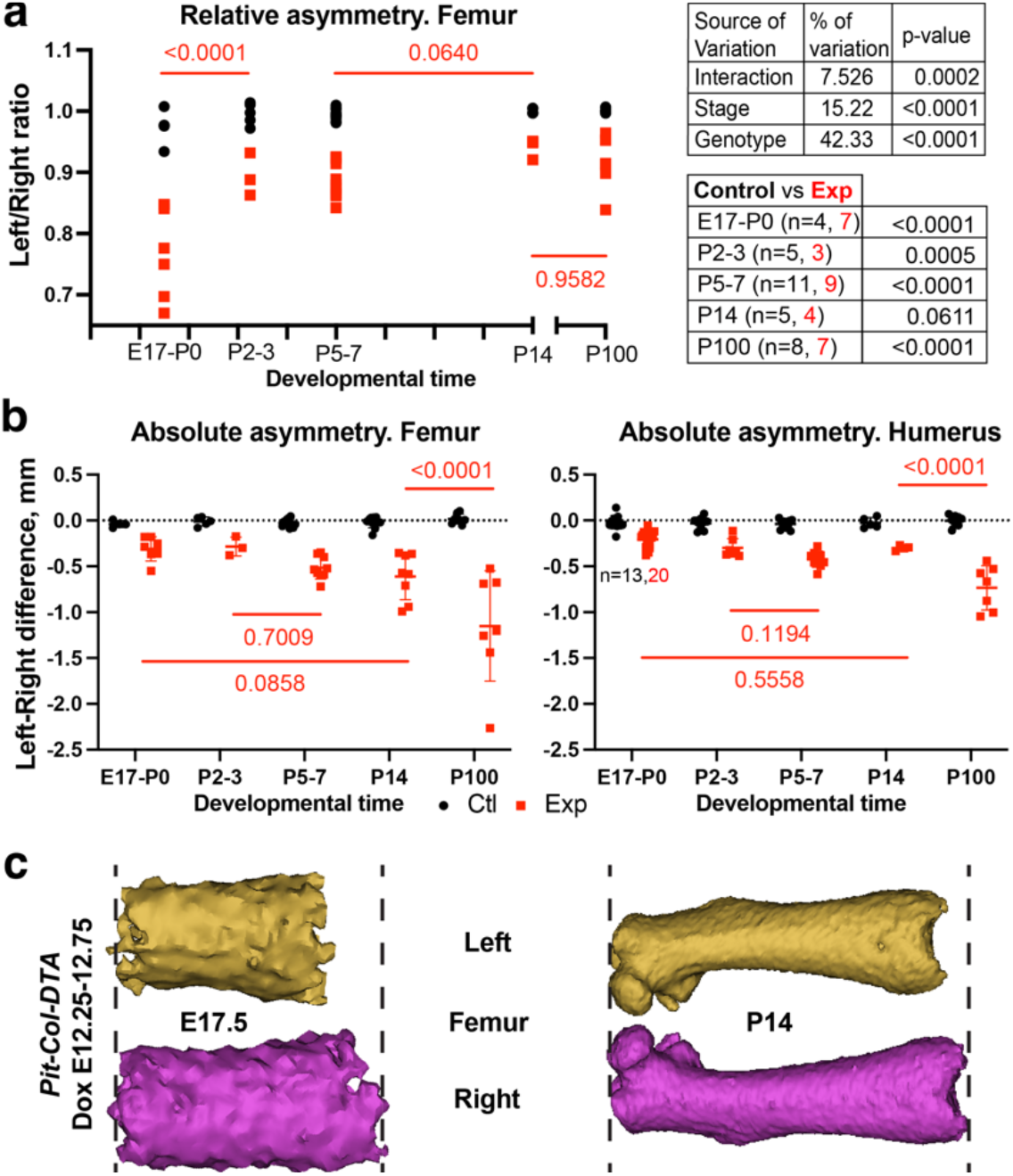
After transient cell death, left-right limb asymmetry is followed by relative catch-up growth. **a** Relative leftright difference (normalised over the right bone size) for control (black) and *Pit-Col-DTA* specimens (red) at the indicated stages. Values for 2-way ANOVA are shown in the Table on the top right. p-values of Sidak’s multiple comparisons tests shown on the graph and the bottom right table. **b** Like **a**, but showing absolute asymmetry values. *n* as in **a**, except where indicated. **c** 3D models of *Pit-Col-DTA* E17.5 and P14 experimental femora, reconstructed from micro-CT images. The models have been re-scaled to show a similar length of the shaft, to allow for comparison of the asymmetry at equivalent scales.

In growing limbs, the recovery of relative bilateral symmetry can have two causes: 1) a roughly constant absolute left-right (L-R) difference becomes relatively smaller as the animal gets bigger; 2) the generation of bone length at a faster rate in the injured limb than in the contralateral one, so that the absolute L-R difference becomes smaller. To determine which aspect(s) is/are contributing to the observed recovery, we plotted the absolute L-R difference of bone length over time. These measurements showed that, in the first two weeks after birth, the L-R difference does not change significantly (Fig. 4b). Eventually, the absolute L-R difference did worsen (Fig. 4b, P100), but not proportionally to the increase in limb size, so that the L/R ratio did not change much as compared to P14 (Fig. 4a-c, Ext. Data Fig. 3). Fig. 4c exemplifies how a similar or bigger absolute L-R difference is relatively minor at P14, as compared to being very evident at E17.5.

### Compensatory cartilage growth is driven by shifts in the proliferation-differentiation balance of chondrocyte progenitors, and by exacerbated cellular hypertrophy

The results above suggest that, in response to chondrocyte loss, the spared cells exhibit compensatory behaviours that repair the damage to the cartilage and limit its impact on bone growth. First, to determine whether compensatory proliferation was taking place (akin to that described by one of us. in a mosaic model of cartilage cell-cycle arrest^19^), we quantified the incorporation of the thymidine analogue 5-Ethynyl-2-deoxyuridine (EdU), provided 1.5 hours before collection. Interestingly, this assay did not reveal differences between Exp L and either Exp R or Ctl limbs at E17.5 or P0 (Ext. Data Fig. 4), when the recovery process should be taking place.

We next reasoned that even if the instantaneous proliferation rate was not significantly changed, it was possible for the flux of resting to proliferative to hypertrophic chondrocytes to change in response to injury. To test this possibility, we performed pulse-chase experiments. We provided a pulse of EdU at either E15.5, E17.5, P1, or P3, and analysed the distribution of EdU^+^ cells two days later, i.e., at E17.5, P0, P3, or P5 (Fig. 5a). In the E15.5→ E17.5 experiment, we observed more EdU^+^ cells in the Exp L RZ as compared to Ctl, and fewer in the Exp L PZ as compared to Exp R (Fig. 5b left and middle). This was independently confirmed by measuring cell density in the E15.5, E17.5 and P0 RZ, which showed progressive increase in Exp L (Ext. Data Fig. 5a). To assess the speed of proliferation in the PZ, we quantified the number of columns with ≥4 EdU^+^ chondrocytes (C4+ hereafter), because two or more cell divisions were expected to take place in two days^60^. Fewer C4+ were found in the Exp L as compared to the Exp R and Ctl limbs (Fig. 5b right). Overall, this early response, together with the cell loss due to cell death, likely explains why the height of the Exp L cartilage is reduced at E18.5 (Fig. 3e). Two days after an E17.5 pulse, the Exp L showed slightly increased number of EdU^+^ cells in the RZ as compared to Exp R, but not as compared to Ctl (Fig. 5c left), and reduced number of C4+, as compared to the Exp R and Ctl (Fig. 5c right). In the P1→P3 pulse-chase, EdU^+^ cells were more abundant in Exp L vs. Exp R across the PZ, and more C4+ were found in Exp L PZ (Fig. 5d, e). Lastly, the P1→P3 pulse-chase revealed increased flux towards the Exp L HZ (not shown), which was even more noticeable in the P3→P5 experiment (Fig. 5f). In summary, in the first step of the response label-retaining progenitors were accumulated in the RZ, while in the postnatal response there was an accelerated transition towards the subsequent states.

**Fig. 5.**
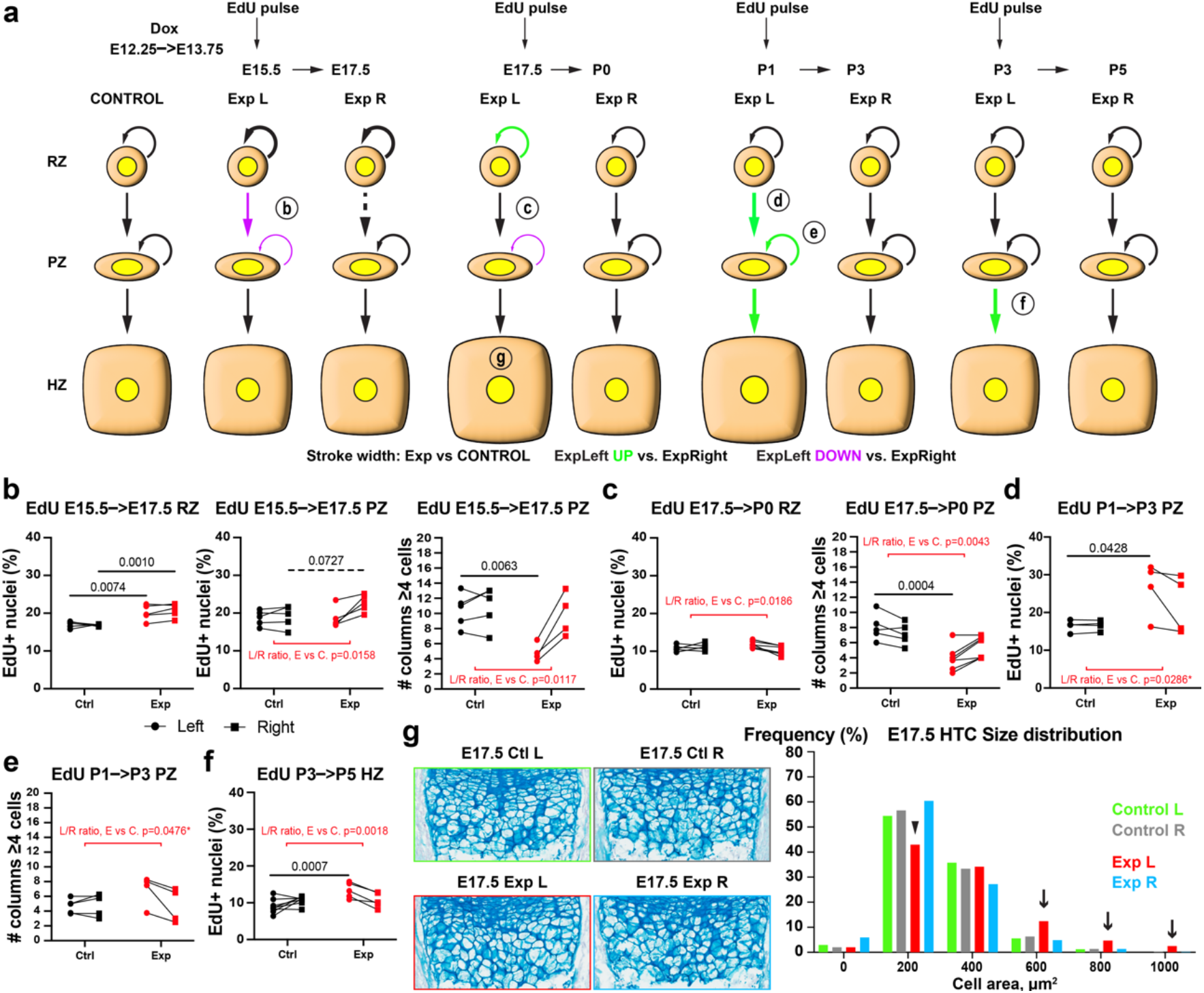
Cartilage repair is driven by a proliferation-differentiation shift in chondrocyte progenitors, and by exacerbated cellular hypertrophy. **a** Schematic summary of the EdU pulse-chase experiments during and after transient unilateral cell death in the cartilage. The stroke width is used to compare Exp L vs. Ctl, whereas the colour is used to compare Exp L vs. Exp R, as indicated. Dashed lines indicate a borderline trend. The main changes are labelled with a circled letter (**b** to **g**), corresponding to the panel where the supporting evidence is presented. **b**-**f** Graphs showing the number of EdU^+^ nuclei and the number of chondrocyte columns with ≥4 EdU^+^ cells, at E17.5 (**b**, n=5 Ctl, 7 Exp), P0 (**c**, n=4,4), P3 (**d, e**, n=5,4) and P5 (**f**, n=8,5), in the indicated cartilage regions (RZ/PZ/HZ, resting/proliferative/hypertrophic). p-values for post-hoc multiple comparisons tests after 2-way ANOVA are shown in black. t-test of Exp L/R ratio vs. Ctl L/R ratio are shown in red (asterisk: non-parametric test was used for non-normal data with few samples). **g** Left: Cell size in the HZ is revealed by alcian blue staining of the extracellular matrix. Right: distribution of HTC sizes in the proximal tibia, for the indicated genotypes and sides, at E17.5 (n=3 Ctl and 3 Exp). Genotypes and sides are colour-coded.

It is noteworthy that, despite the decreased number of cells being produced during the injury phase, the length of the HZ was never much affected (Fig. 3e), suggesting the participation of other compensatory mechanisms. We thus measured HTC cell area on histological sections at multiple stages, finding that the size distribution in the Exp L HZ shifted away from small areas (Fig. 5g arrowhead) and towards high ones (Fig. 5g arrows), including values that were never found in Exp R or Ctl HZ. This compensatory hypertrophy only lasted for a few days, as the size distribution returned to being mostly normal by P3 (Ext. Data Fig. 5b).

The retention of EdU^+^ cells in the RZ during the early phase of the recovery (Fig. 5b) resembled the enhanced self-renewal of Axin2^+^ chondroprogenitors observed during transient diet restriction in juvenile mice^1^. Since the capability of self-renewal in chondroprogenitors has been shown to appear only at postnatal stages^32^, one possible explanation was that self-renewal had been precociously activated upon injury. The alternative scenario was that there was no enhanced production of RZ chondrocytes, but rather retention without proliferation. Importantly, the fact that the left/right ratio of instantaneous EdU incorporation increased in Exp animals between E15.5 and E17.5 (Ext. Data Fig. 6) did not support the second scenario, suggesting indeed increased self-renewal. However, we were not able to test whether chondroprogenitor self-renewal was taking place, as the expression of the typical markers of <u>postnatal</u> long-term chondroprogenitors/cartilage stem cells such as Axin2, FoxA2 and PTHrP^34,61,62^, was undetectable by immunofluorescence (not shown). Moreover, bulk RNA-seq of the growth plate did not reveal major changes in the otherwise low levels of these markers, except for a non-significant trend towards higher expression of *Pthlh*, the gene coding for PTHrP, in Exp L cartilage at E15.5 and P0 (Ext. Data Fig. 7a-b).

**Fig. 6.**
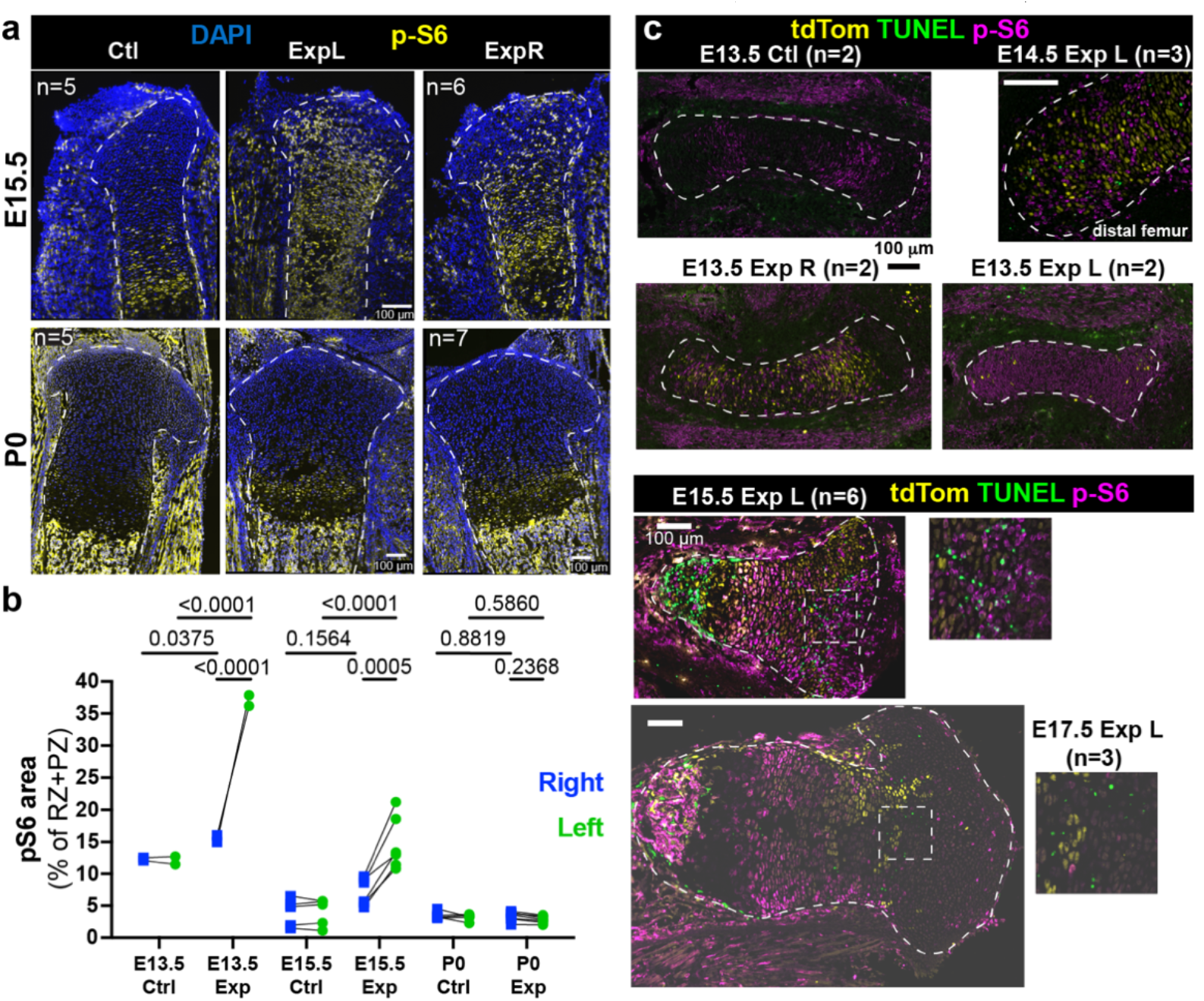
Ectopic mTORC1 activity after cell death in the cartilage. **a** p-S6 signal reveals mTORC1 activity in Ctl and Exp groups. Sample number (n) as indicated. **b** p-S6^+^ area, as % of the RZ + PZ (n= 2, 2, 5, 6, 5, 7, from left to right). **c** Time course of p-S6 immunostaining in cartilage, and close-ups to show the spatial relationship with cell death (at E14.5 in the distal femur and E13.5 and 15.5 in the proximal tibia). n as indicated in the panels.

### The mTORC1 pathway initiates the compensatory response in the injured cartilage

In order to uncover the molecular mechanisms that mediate the cellular behaviours described above, we performed bulk RNA-seq on left and right cartilage (from femur and tibia) of foetal and early postnatal mice (see Methods and Ext. Data Fig. 7a). To reveal candidate genes/pathways involved both in the early response to injury and in the subsequent recovery, we performed Differential Gene Expression (DGE) analysis and Gene Set Enrichment Analysis (GSEA) between Exp L, Exp R and Ctl at E15.5, E17.5, P0 and P3 (Ext. Data Fig. 7c-d and Ext. Data Tables 1-4). While several enriched pathways called our attention, assessment of candidate expression confirmed only a few of them as differentially expressed. Namely, we immuno-stained for phosphorylated ribosomal protein S6 (p-S6) as an mTORC1 activity readout, and found it ectopically expressed in the Exp L cartilage during the injury phase, going back to a normal state by P0 (Fig. 6a-c). Surprisingly, the Exp R cartilage also showed increased mTORC1 signalling as compared to Ctl cartilage at E15.5, although to a lesser extent than Exp L (see Discussion section). A time-course of p-S6 immunostaining revealed that, at E13.5, p-S6 expression was widespread in the Exp L cartilage, (Fig. 6c), despite the fact that TUNEL signal only became detectable later (E14.5), and in a patchy manner only. This suggests that the mosaic expression of DTA triggers a non-autonomous stress response in most of the cartilage, eventually killing some of the chondrocytes cell-autonomously. At E15.5, the p-S6 signal was patchier, and closely associated with the dying cells (Fig. 6c insets), suggesting that cell death triggers a short-range response that maintains mTORC1 activity.

To determine the relevance of the ectopic expression of mTORC1, we set out to inhibit mTORC1 activity *in vivo*, during the prenatal period post-injury. *Pitx2-Cre; Col2a1-rtTA* females time-mated with *Tigre*^*Dragon-DTA*^ males were provided with Dox from E12.25 to 13.75, and then with the mTORC1 inhibitor Rapamycin (4mg/kg twice daily) from E14.25 to E18.75 inclusive (Fig. 7a). While the treatment did not worsen the L/R asymmetry of experimental animals by P7, likely due to incomplete inhibition of mTORC1 activity by the *in utero* treatment (Extended Data Fig. 8a, b), it was however enough to impair the subsequent recovery of cartilage cytoarchitecture. In contrast to untreated litters, in Rapamycin-treated litters the left/right ratio of the length of each cartilage region was significantly smaller in experimental (rtTA^+^) animals, as compared to controls (Fig. 7b, b’, red symbols). Interestingly, Rapamycin seemed to affect both control and experimental cartilage, especially the proliferative and resting zones. As a consequence, the internal proportions (i.e. the stratification) of the control cartilage changed, with the RZ being significantly longer–and the PZ shorter–in treated animals as compared to untreated ones (Fig. 7c). Besides this general effect, however, Rapamycin also seemed to inhibit the compensatory response in the Exp L cartilage. Indeed, the Exp L cartilage was shorter than the Exp R at P7, both at the RZ and PZ levels (Fig. 7d), whereas in untreated Exp animals all growth plate zones were similar between the left and right limbs (Fig. 3e). Along these lines, while the effect of Rapa on controls is towards increasing the length of the RZ (Ext. Data Fig. 8c), in Exp L this effect is counterbalanced by the failure of the compensatory response when mTORC1 is inhibited. Lastly, we measured the SOC in the P7 Rapa-treated samples, and found that the Exp L/R ratio was not significantly different from that of Control animals (Fig. 7e), unlike in untreated samples (see Ext. Data Fig. 2). These results indicated that the delay in Exp cartilage progression also requires mTORC1 activity.

**Fig. 7.**
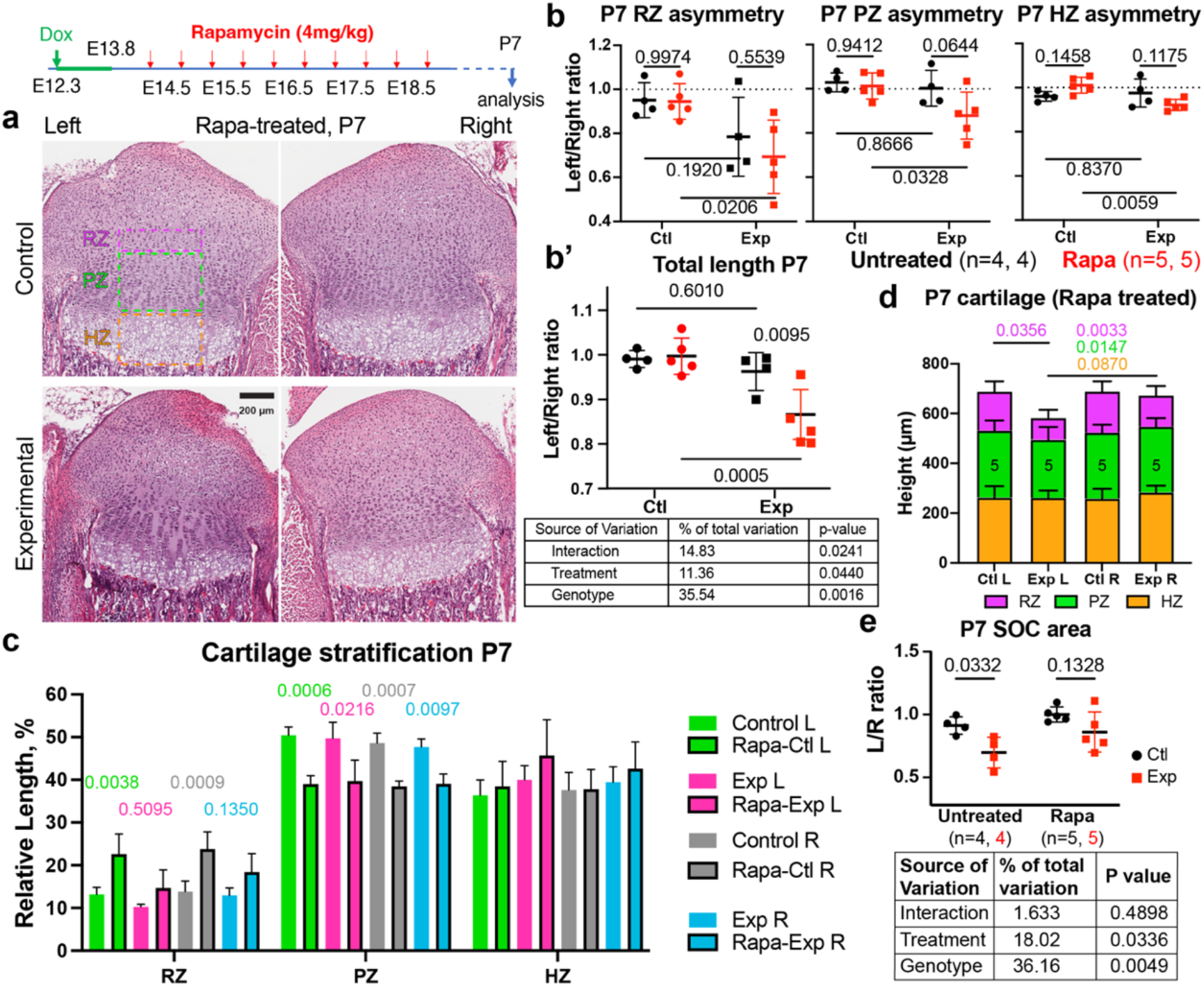
mTORC1 signalling is required to repair the damaged cartilage. **a, b** Pregnant females were treated with the mTORC1 inhibitor Rapa as indicated, and pups collected at P7 for analysis. H&E images (**a**) and the average of the longest length per section (**b, b’**) are shown as L/R ratios for Ctl and Exp samples, treated or not with Rapa. RZ/PZ/HZ, resting/proliferative/hypertrophic zones. Dotted lines: expected L/R ratio for controls. The table shows the results of the 2-way ANOVA. p-values for the multiple comparisons tests are shown on the graphs. **c** Relative length of the different proximal tibia zones in P7 animals from untreated litters or treated *in utero* with Rapa. p-values for the multiple comparisons tests (after pairwise 2-way ANOVAs) are shown. **d** Like **c**, but only for treated litters. Note the differences between Exp L and Exp R. **e** L/R ratio of the secondary ossification centre area in untreated and Rapa treated P7 samples. 2-way ANOVA and Sidak’s tests are shown

## Discussion

### Catch-up growth as a tool to uncover the mechanisms controlling organ size and repair

Developmental robustness is a biological phenomenon that can not only inspire regenerative therapies but also reveal mechanisms that regulate organ growth. In this study we set out to explore the molecular and cellular mechanisms behind catch-up growth of the long bones. This is a unique organ model that grows via a cartilage template that is produced and destroyed in opposite sites at the same time. This process requires tight control of the proliferation and differentiation of cartilage progenitors, and leads to layers of different chondrocyte states being organised in a distinct cytoarchitecture. The proportions of this layered structure often change across different growth plates, developmental stages and species, and have been shown to correlate with the speed and/or extent of long bone growth^63,64^. However, how these proportions are established and/or recovered after a perturbation is not completely understood, and was explored in this study.

### A new mouse model for acute cell death in the growth plate cartilage

We capitalised on the *Dragon-DTA* allele that we described previously^38^ to develop a model of acute unilateral injury in the cartilage. A brief pulse of Dox was enough to induce transient expression of DTA and subsequent cell death in the foetal cartilage, mostly in the left limb (Fig. 2). Unsurprisingly, the extensive cell death that takes place in the Exp L cartilage caused major disruptions in its integrity, with acellular gaps that in some cases extended across almost the whole growth plate (Fig. 3). As a consequence, the length of the growth plate was altered, especially in the case of the RZ and PZ. Given that the growth plate is the template for bone growth, this model also showed a remarkable growth impairment of the left-limb bones, causing a peak left-right difference of 25-30% by P0 (Fig. 4). Also by P0, cell death was over, potentially allowing for recovery to happen. This model thus provides at the same time exquisite tissue specificity and control of the timing and duration of the injury. It is also versatile, as it can be combined with any Cre and (r)tTA lines to affect virtually every tissue of interest, which could be very valuable for developmental and regenerative biologists.

### Recovery of cartilage integrity and cytoarchitecture after cell death is in part mediated by mTORC1 activity

After the injury, we observed a remarkable recovery of cartilage integrity and cytoarchitecture (Fig. 3), associated with a biphasic response (Fig. 5): first, increased retention of EdU^+^ chondrocytes in the resting zone between E15.5 and E17.5, at the expense of the transition to the proliferative pool; second, a reversal of this balance, i.e. accelerated transition towards first the proliferative and then the hypertrophic pools between P1 and P5. During the first phase, compensatory hypertrophy also took place in the HZ, explaining why its size was not affected despite the reduced cell output from the upstream layers. Based on the literature and RNA-seq data, we hypothesised that these behaviours could be associated with increased mTORC1 activity, which was confirmed at the histological level by assessing expression of p-S6 (Fig. 6). Interestingly, p-S6 has been recently associated with the injury response area in different types of skin wounds^35^, muscle injuries^65^ and limb amputation, suggesting that it is a conserved response. In the case of skin wounds, p-S6 was shown to be a modulator of the response, but not strictly necessary for the response to happen^35^. It remains to be determined whether this is also the case in our model of catch-up growth, as we could not achieve full *in vivo* inhibition. Partial inhibition did however impair the recovery of cartilage cytoarchitecture in our model (Fig. 7), suggesting it is quite important. A priori, this role of mTORC1 could seem at odds with the findings by Oichi et al. ^1^, as in their model of catch-up growth they showed that the bias towards self-renewal at the expense of transition to the proliferative pool was associated with decreased expression of *Igf1* in the resting zone, which could presumably result in reduced mTORC1 signalling. However, mTORC1 activity was not assessed in their study, leaving open the possibility for it to be active, and thus independent of IGF1 in the RZ. Alternatively, mTORC1 may only be required in the response to physical damage (our study), but not in diet restriction, or only during foetal stages (our study) and not in juvenile mice. Additional studies will help elucidate these questions.

### A new hybrid model of catch-up growth

As a consequence of impairing Exp L cartilage function and integrity, our unilateral injury model also caused a remarkable asymmetry in bone length, with the left femur being ∼25% shorter by E17.5-P0 (Fig. 4). In terms of catch-up growth, while the relative asymmetry became milder with time, suggesting functional recovery, the absolute left-right difference did not improve over the whole growth period. In order to classify this behaviour as catch-up growth or not, it is important to consider that, in a growing animal, triggering cell death in the cartilage will also ablate the long-lived chondroprogenitors. Therefore, this is expected to not only reduce the amount of new bone produced concomitantly to (and shortly after) cell death, but also to impair the subsequent growth potential, due to reduction of the pool of progenitors. In summary, in the absence of compensatory mechanisms, the absolute left-right difference is expected to continuously grow over time (Fig. 8a). On the other side of the spectrum, if there are compensatory mechanisms that can fully replenish the pool of progenitors, such that there is no loss of potential, even if it involves temporal stalling, then absolute asymmetry should decrease over time, as long as there is enough time to complete the growth period (Fig. 8b). In the former case, the growth curve shifts vertically, whereas in the latter, the growth curve shifts horizontally. What we think we have found with our model is a hybrid situation, in which part of the potential is lost during the injury, but the remaining potential is preserved during the stalling phase, such that when growth resumes, the growth curve shifts diagonally (Fig. 8c). Interestingly, this would lead to the absolute left-right difference being roughly maintained over the linear growth period, which is what we observed (Fig. 4). Of note, the preservation of growth potential that we postulate for this model would be the first case in which this behaviour has been observed during foetal stages, before the radical switch in clonal behaviour that happens postnatally^32^. It remains to be determined whether all chondroprogenitors are capable of this behaviour, or just a sub-population of them.

**Fig. 8.**
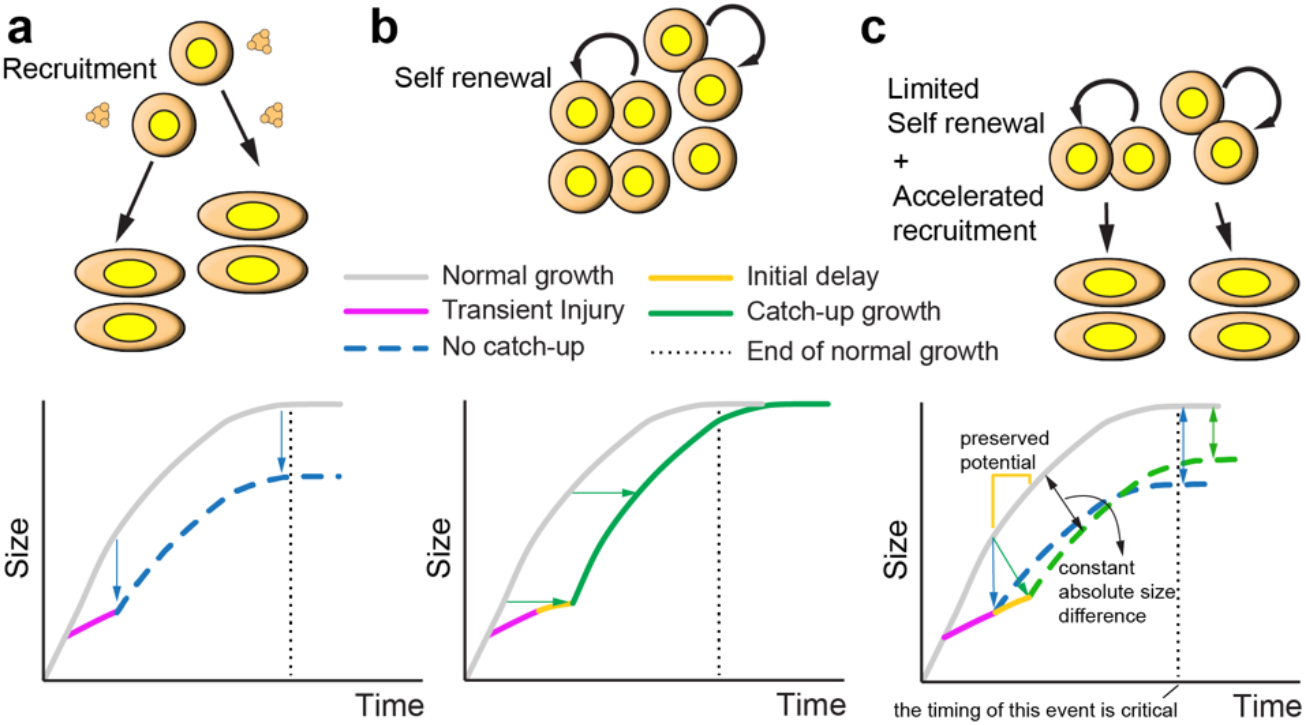
A new hybrid model of catch-up growth. **a-b** The two expected potential responses to cell ablation. In the absence of self-renewal after injury (a), the chondroprogenitors get recruited into the proliferative pool, using up their growth potential, which shifts the growth curve vertically, leading to absence of catch-up growth. In the case of perfect self-renewal (b), no potential is lost, and when growth resumes, the growth curve is shifted horizontally, so that catch-up growth happens over time. **c** The new hybrid model, in which a delay phase after the injury freezes up the growth potential, so that when growth resumes, it does it at an intermediate speed. Normal size is never recovered, but the deficiency does not become worse. Note the improvement with respect to the absence of catch-up.

Importantly, to maximize the catch-up in these models (8b and 8c), it is necessary that the growth period is extended beyond the normal time, which is why we analysed our model at P100, instead of at 9 weeks of age, (i.e., when normal longitudinal growth ceases in mice). It follows that if there is a systemic growth halt before a full local recovery has happened, catch-up growth will be interrupted, which is something we cannot discard in our study. Future studies will need to investigate whether this is the case and, if so, how to extend the growth period beyond the normal time, to improve the efficacy of growth therapies.

### The control of normal cartilage proportions by mTORC1

Besides the role of mTORC1 in the compensatory response, our study has also revealed a potential role in the maintenance of the layered structure of the cartilage during normal growth. Indeed, treatment of control animals with Rapamycin led to changes in the relative proportions of RZ, PZ and HZ (Fig. 7). Given how variable cartilage stratification is across developmental stages, different bones and different species, further exploring the role of mTORC1 in coordinating the size of the growth plate zones is warranted. We speculate that it could be mediated via its known interactions with IHH^66^ and PTHrP^33^, and other interactions yet to be determined.

### A potential systemic response triggered by the unilateral injury

During characterisation of the injury model, we realised that mTORC1 activity was also ectopically activated in the Exp R cartilage, especially after E15.5 (Fig. 6). While at first this may seem like a direct effect of leaky DTA expression in the right cartilage, several lines of evidence go against this possibility. Namely, the timing and pattern of the activation of mTORC1 are quite different between Exp L and Exp R cartilage. Indeed, while p-S6 was detected all over the Exp L cartilage as early as E13.5 (Fig. 6), its expression was not obvious until E15.5 in the Exp R cartilage, when expression in Exp L had already switched to a salt & pepper pattern. These results suggest that there are two waves of ectopic mTORC1 activity in the Exp L cartilage, and that the Exp R only experiences the second one. We hypothesise that the cartilage injury is triggering not only a local response but also a systemic one, which includes activation of mTORC1 at distant sites. There are several examples in the literature where this has been observed. For example, unilateral injury in the limb muscles in adult mice leads to activation of mTORC1 signalling and of the so called G_0_ alert state in muscle stem cells of not only the injured limb, but of the uninjured one as well^65^. Similarly, unilateral limb amputation in axolotl leads to systemic activation of stem cells, including increased mTORC1 signalling and additional rounds of cell replication^67^. Interestingly, in our model we also detected cell behaviour changes in the Exp R chondrocytes, such as the increase retention of EdU^+^ chondrocytes in the resting zone between E15.5 and E17.5, despite the absence of a major injury (Fig. 5). Importantly, we would not have been able to detect this contralateral effect with traditional approaches, as foetal mice are quite inaccessible for fine manipulation of a single limb tissue, while traditional genetic injury models typically affect left and right limbs equally. From an evolutionary perspective, this type of systemic ‘priming’ could prepare other regions of the body to respond faster, providing adaptive value in the case that a second injury takes place. Indeed, in the two studies mentioned above, the primed cells were able to respond faster to secondary injuries^65,67^. In future studies, it would be interesting to determine if this is also the case in our model of compensatory growth, and whether this response extends to other organs.

## Author Contributions

C.H.H.: data acquisition and analysis, supervision, figure preparation and funding sourcing. S.A.: data analysis, manuscript editing, supervision, figure preparation. H.C.: data acquisition and analysis, figure preparation. B.Z: data acquisition, figure preparation. D.P.: tool generation, data analysis, supervision. A.R-D.: conceptualization, data acquisition and analysis, funding sourcing, supervision, figure preparation, manuscript drafting.

## Acknowledgments

We thank Jonathan Gleadle, Vincent Wong and Darling Rojas-Canales for inspiring discussions about mTORC1 and the balance between proliferation and cell size in compensatory responses. We also acknowledge the Monash Bioinformatics Platform (especially Kirill Tsyganov), and Trevor Wilson, at the Medical Genomics Facility, Monash Health Translation Precinct, for their excellent technical help.

## Extended Data Figures

Follow the link for Supplemental Information

## Extended Data Tables 1-4

These Tables present the Differential expression (DE) analysis for the RNA-seq experiment described in Ext. Data Fig. 7. Data from E15.5, E17.5, P0 and P3 are presented, respectively in Tables 1-4. Within each table, multiple comparisons are presented in different tabs. C, control; E_L, Exp Left, E_R, Exp Right.

## Data Availability Statement

The RNA-seq datasets generated and analysed for this study can be found in the GEO repository (GSE235779).

