## Supplemental Information for "Compensatory growth and recovery of tissue cytoarchitecture after transient cartilage-specific cell death in foetal mouse limbs"

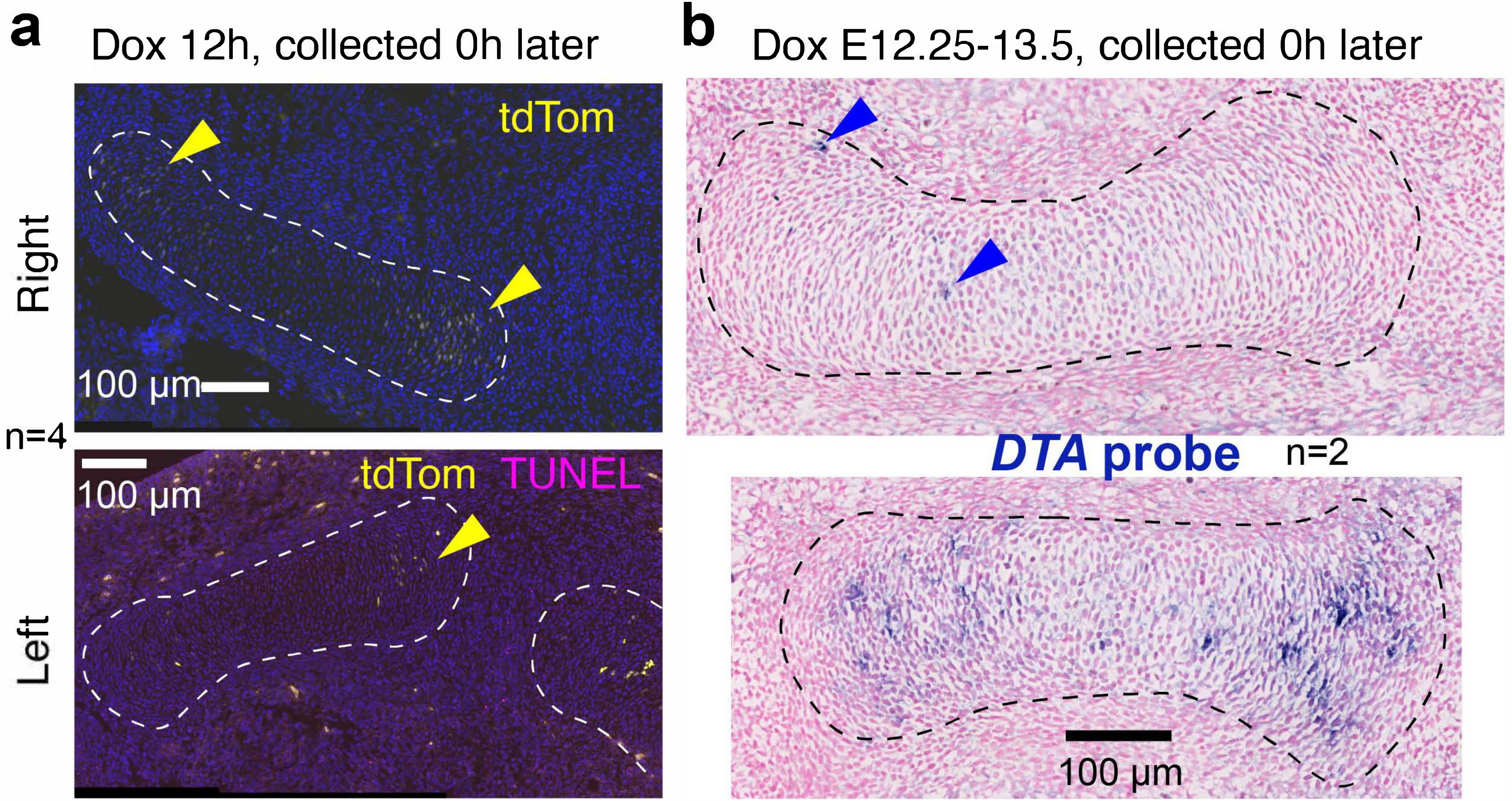


**Ext. Data Fig. 1**. **a** Immunohistochemistry showing the expression of tdTomato (yellow arrowheads) achieved in the cartilage of the *Pit-Col-DTA* model (white dashed lines) after 12h Dox treatment. **b** *In situ* hybridisation showing the expression of *DTA* (blue arrowheads) achieved in the cartilage of the *Pit-Col-DTA* model (black dashed lines) after 30h Dox treatment. Sample number (n) as indicated.

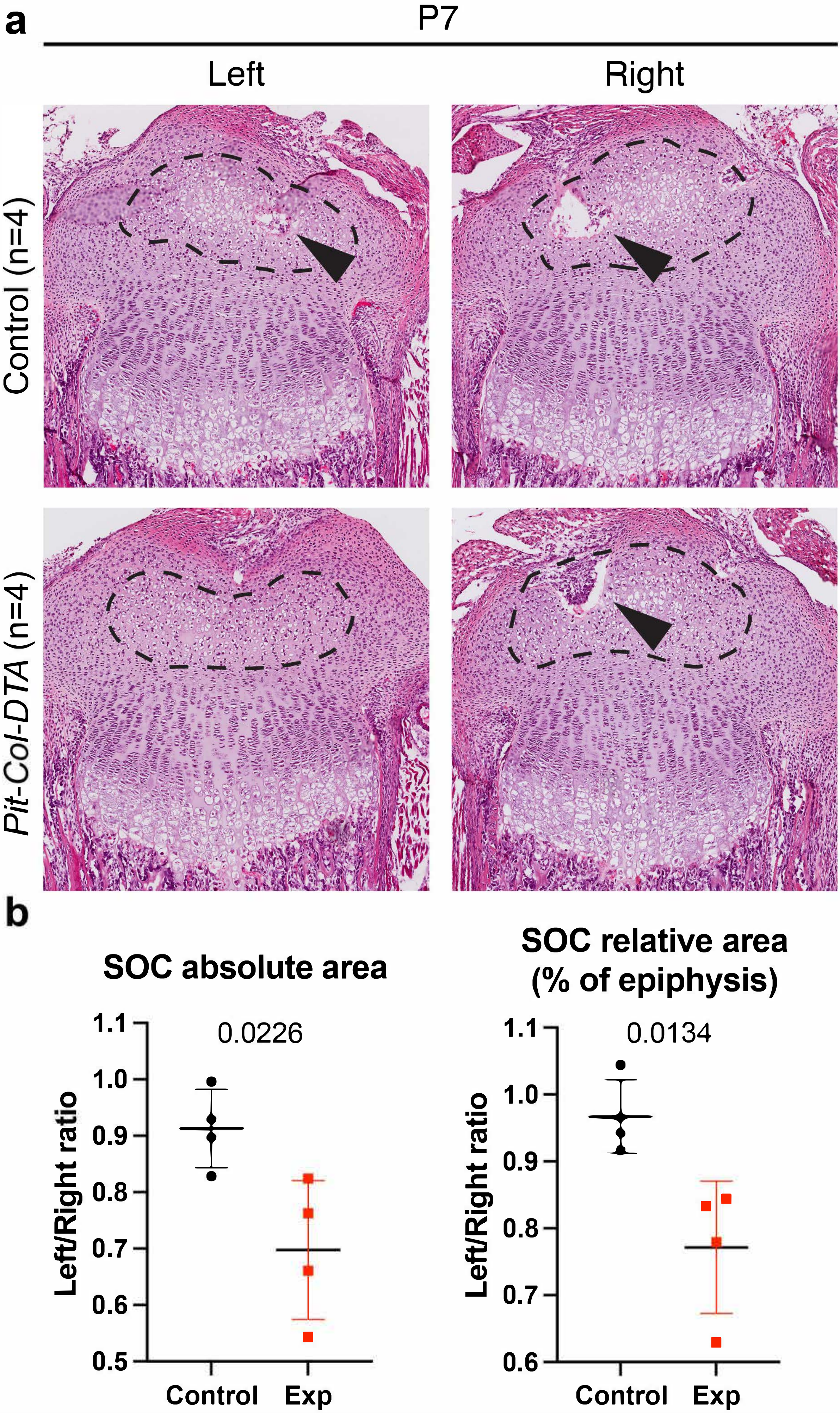


**Ext. Data Fig. 2**. **a** H&E staining on sections of proximal tibia at P7 showing the formation of the SOC (dashed lines) and the invasion of blood vessels (arrows). **b** Quantification of the left/right ratio of the absolute area ($\mu m^2$) of the SOC and blood vessel invaded region (left) and the relative area of SOC plus invaded region over the epiphysis area, at P7. p-values for unpaired t-tests are shown, n=4 Ctl and 4 Exp.

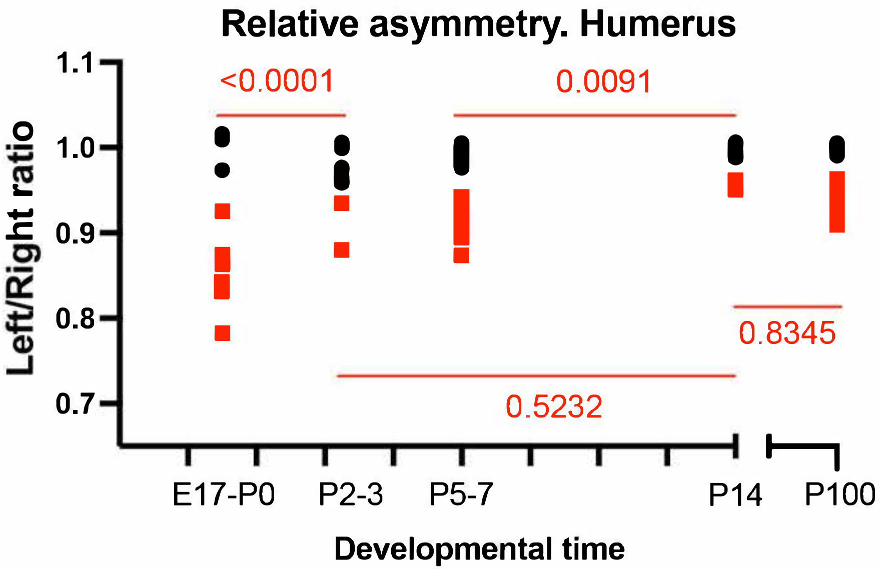


| Source of Variation | % of variation | P value |
|---|---|---|
| Interaction | 10.64 | <0.0001 |
| Stage | 8.574 | <0.0001 |
| Genotype | 47.35 | <0.0001 |

| **Control** vs **Exp** | |
|---|---|
| E17-P0 | <0.0001 |
| P2-3 | 0.0046 |
| P5-7 | <0.0001 |
| P14 | 0.0406 |
| P100 | <0.0001 |

**Ext. Data Fig. 3**. Left/right ratio of the humeri length for control (black) and *Pit-Col-DTA* specimens (red) at E17-P0 (n=4 Ctl, 7 Exp), P2-P3 (n=5,3), P5-P7 (n=11,9), P14 (n=5,4), P100 (n=8,7). Values for 2-way ANOVA are shown in the Table on the top right. p-values of Sidak's multiple comparisons tests are shown on the graph and the bottom right table.

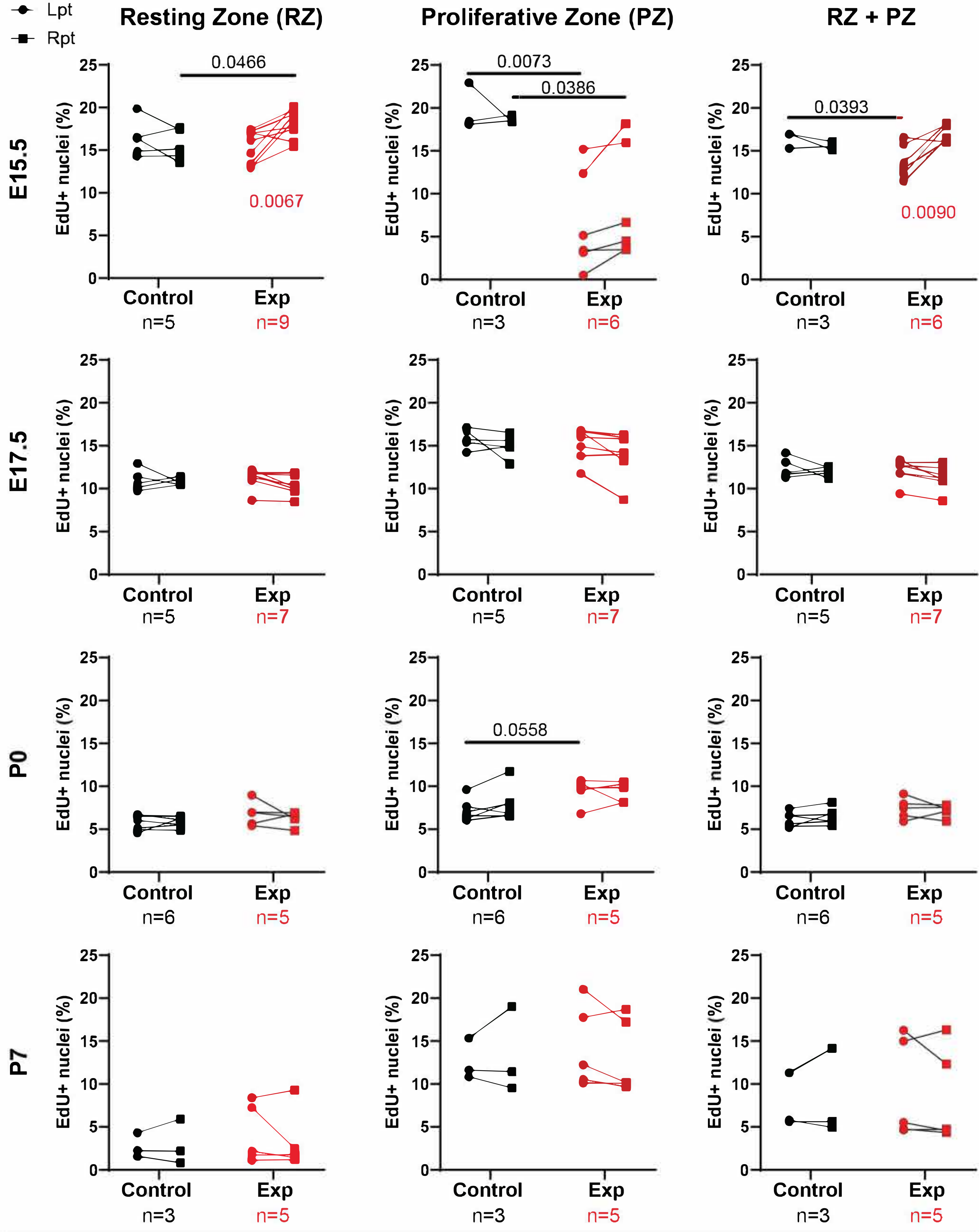


**Ext. Data Fig. 4**. Graphs showing the percentage of EdU+ nuclei/total nuclei of either the RZ, the PZ or both combined, at the indicated stages. Statistical analyses are 2-way ANOVAs and Sidak's post-hoc multiple comparisons tests, performed at each stage for all zones, in a pairwise manner. Sample number (n) as indicated.

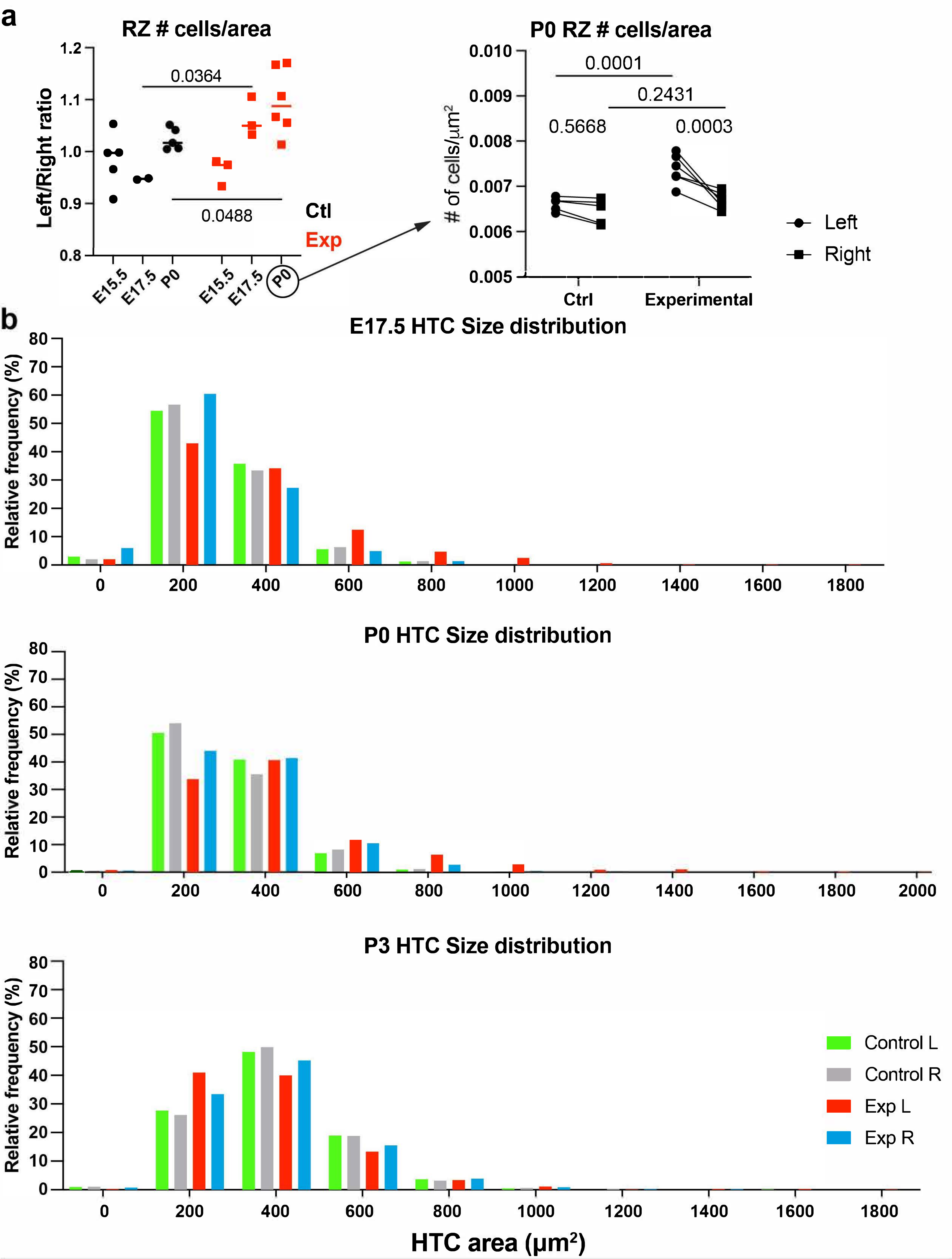


**Ext. Data Fig. 5**. **a Left**, Graphs showing the left/right ratio of the number of cells in the resting zone per area unit at E15.5 (n=5 Ctl, 3 Exp), E17.5 (n=5,3), P0 (n=5,6). Right, Graph showing the number of cells in the resting zone per area unit at P0. **b** Histogram of the cell size frequency distribution of the hypertrophic chondrocytes for Ctl and *Pit-Col-DTA* specimens at E17.5 (n=3 Ctl, 3 Exp), P0 (n=3,3), P3 (n=3,3).

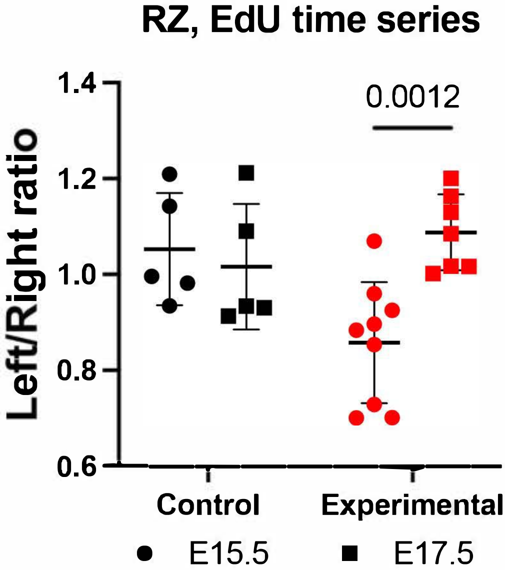


**Ext. Data Fig. 6**. Graphs showing the left/right ratio of the instantaneous EdU incorporation in the resting zone (RZ) chondrocytes for Ctl and *Pit-Col-DTA* mice at E15.5 (n=5 Ctl, 9 Exp) and E17.5 (n=5, 7). Statistical analyses are 2-way ANOVA and p-values of Sidak's multiple comparisons tests shown on the graph

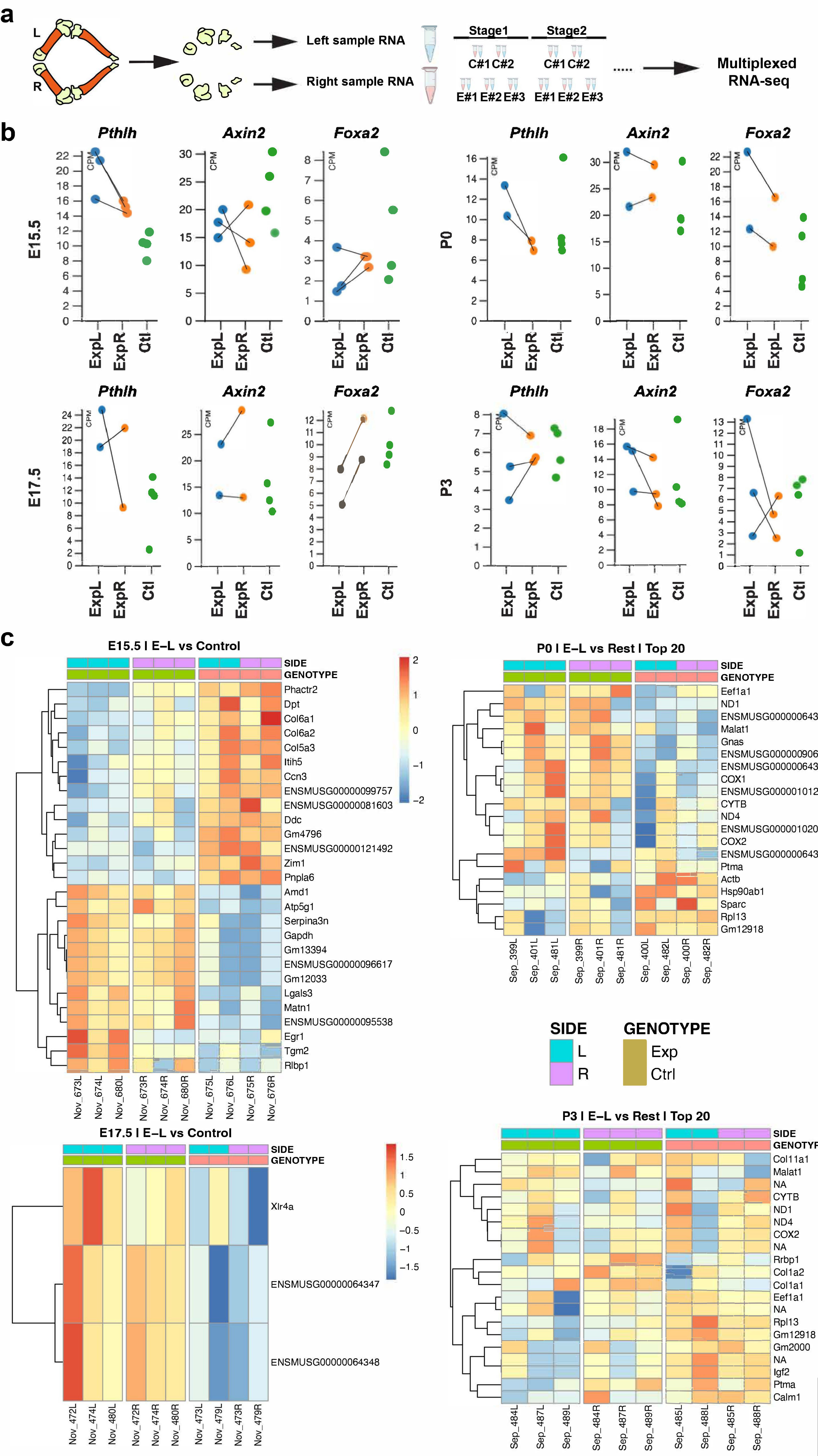


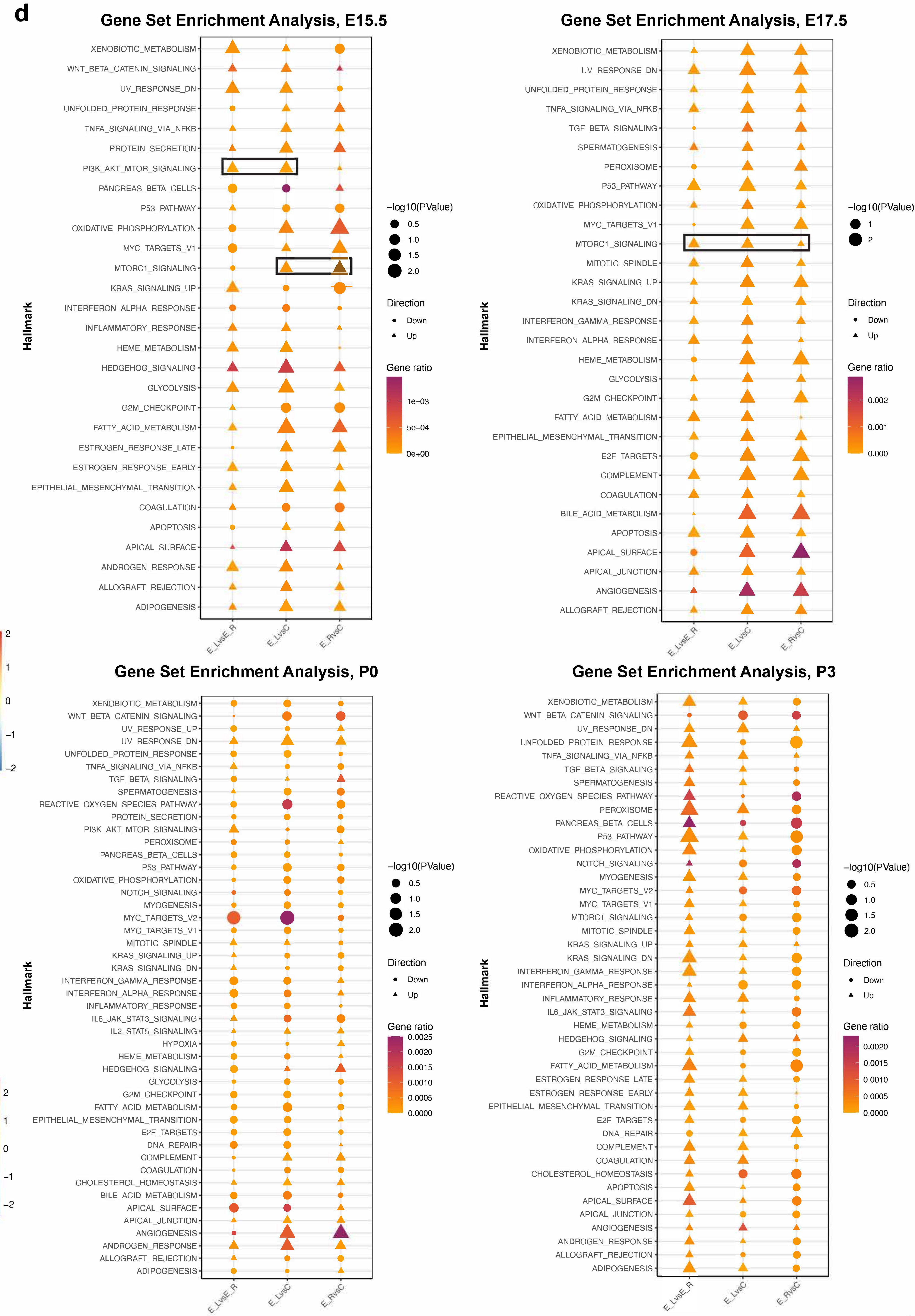


**Ext. Data Fig 7. a** Schematic showing the cartilage region collected and processed for the bulk RNA-sequencing analysis. **b** Expression of *Pthlh*, *Axin2* and *Foxa2*, presented as counts per million (CPM) at E15.5 (n=2 Ctl, 3 Exp), E17.5 (n=2,2), P0 (n=2,2) and P3 (n=2,3), analysed using Degust software. **c, d** Heatmaps of the Differential Gene Expression (DGE, **c**) and Gene Set Enrichment Analyses (GSEA, **d**) at the indicated stages, as described in the Methods. The mTORC1 related hallmarks are outlined.

**a**

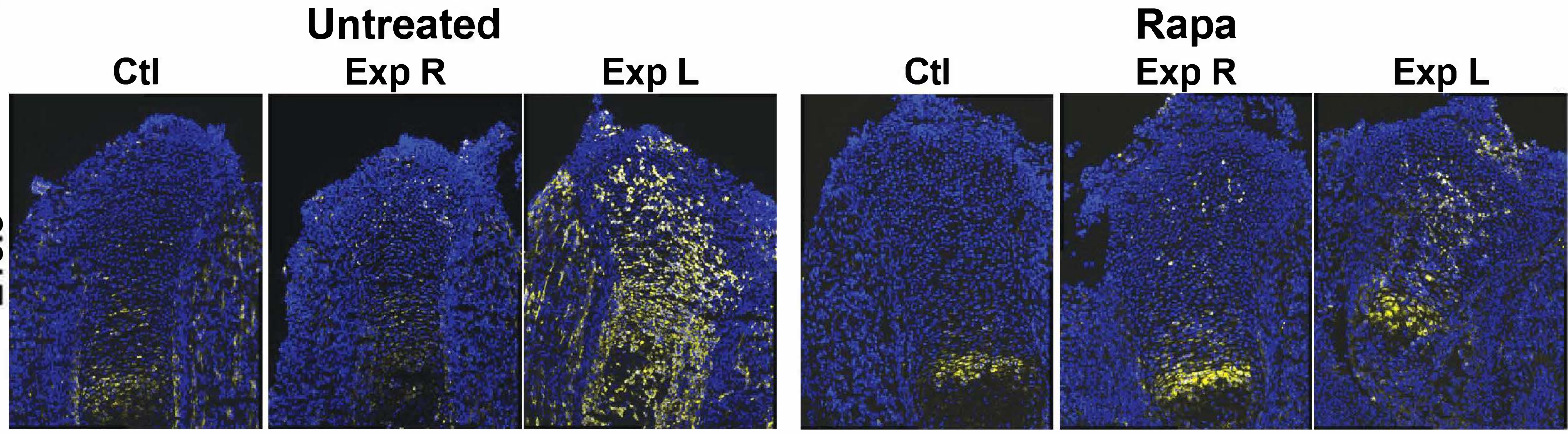


**b**

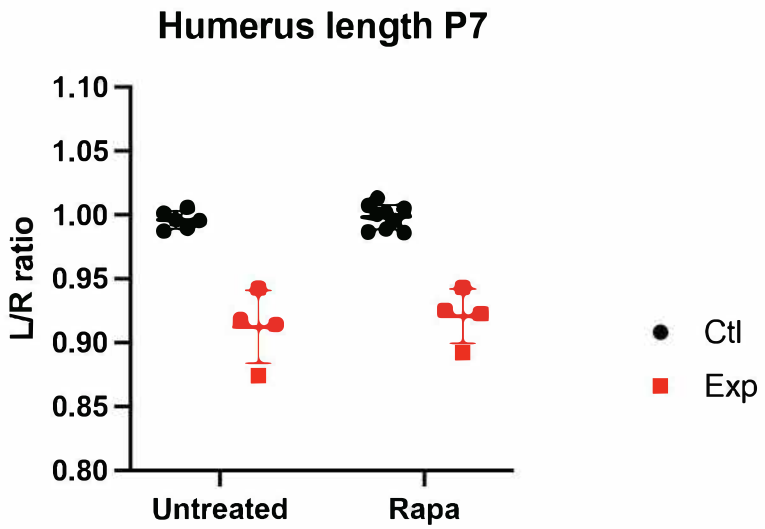


| Source of Variation | % of total variation | P value |
|---|---|---|
| Interaction | 0.1240 | 0.6657 |
| Treatment | 0.3911 | 0.4453 |
| Genotype | 85.59 | <0.0001 |

| Source of Variation | % of total variation | P value |
|---|---|---|
| Interaction | 0.07676 | 0.7488 |
| Treatment | 0.005904 | 0.9292 |
| Genotype | 74.21 | <0.0001 |

**c**

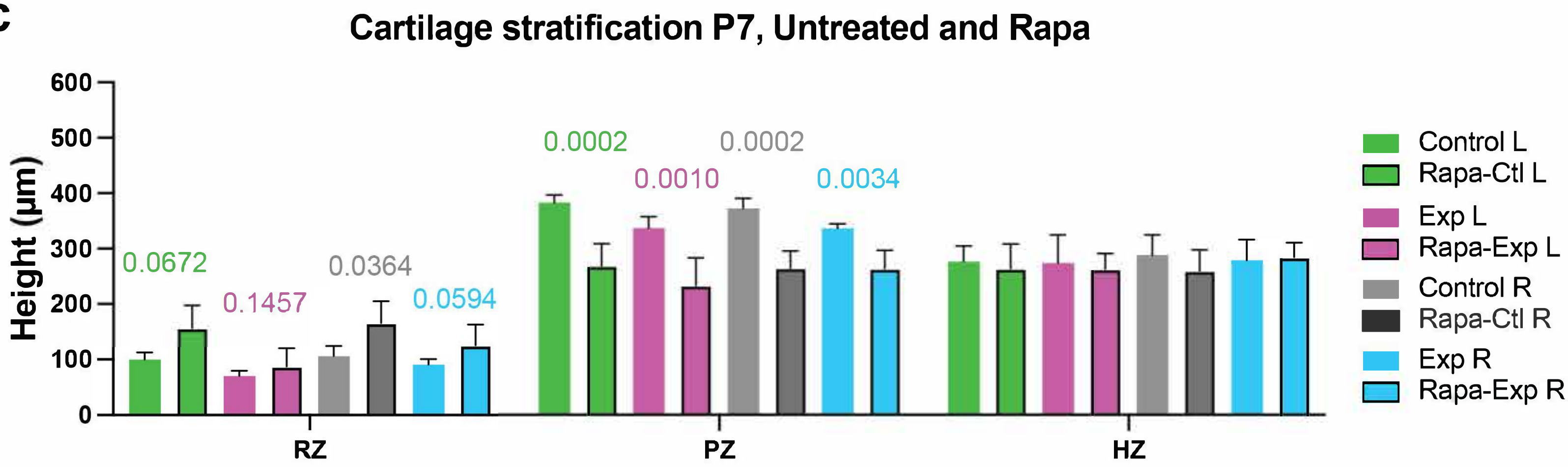


**Ext. Data Fig 8. a** Immunohistochemistry for p-S6 (yellow) in Untreated (n=5 Ctl, 6 *Pit-Col-DTA*) and Rapamycin-treated (Rapa, n=4,6), specimens at E15.5. **b** Graphs showing the left/right ratio of the length of P7 humerus (Untreated, n=6,4; Rapa, n=9,4) and femur (Untreated, n=9,8; Rapa, n=9,4) as indicated. Values for 2-way ANOVA are shown in the Table on the bottom of each graph. **c** Absolute length (µm) of the different cartilage zones (proximal tibia) in P7 animals from untreated litters or treated in utero with Rapa. p-values for the multiple comparisons tests (after pairwise 2-way ANOVAs) are shown.
